# A Minimal Stochastic Model of Microbial Ecological Dynamics in a Single-Species-Single-Resource Setting

**DOI:** 10.64898/2026.07.01.735782

**Authors:** Cheuk Fung Alvin Leung, Anatoly B. Kolomeisky

## Abstract

Microbes exhibit complex dynamic behavior as the result of a large number of biochemical processes, spatial and temporal interactions, environmental variations, and evolutionary pressure. Although significant progress has been achieved in understanding microbial ecological dynamics, multiple open questions remain, including the microscopic mechanisms of growth and the roles of nutrients and stochasticity. In this work, we present a minimal theoretical approach to clarify the link between consumption of resources by microbes and their growth. A stochastic model that accounts for a single microbial species consuming a single type of resource while growing via cell division is studied analytically and via Monte Carlo computer simulations. We identify three distinct dynamical regimes of microbial growth determined by the relative magnitudes of resource uptake and division rates and initial conditions. We also show that stochasticity influences the dynamic behavior when the amounts of microbes or resources are low. The model recovers Monod growth kinetics and provides a mechanistic interpretation of the Monod constant and maximal growth rate. The theoretical framework presented captures a wide spectrum of dynamic behaviors in microbial systems, providing a clearer microscopic picture to explain their underlying complex mechanisms.

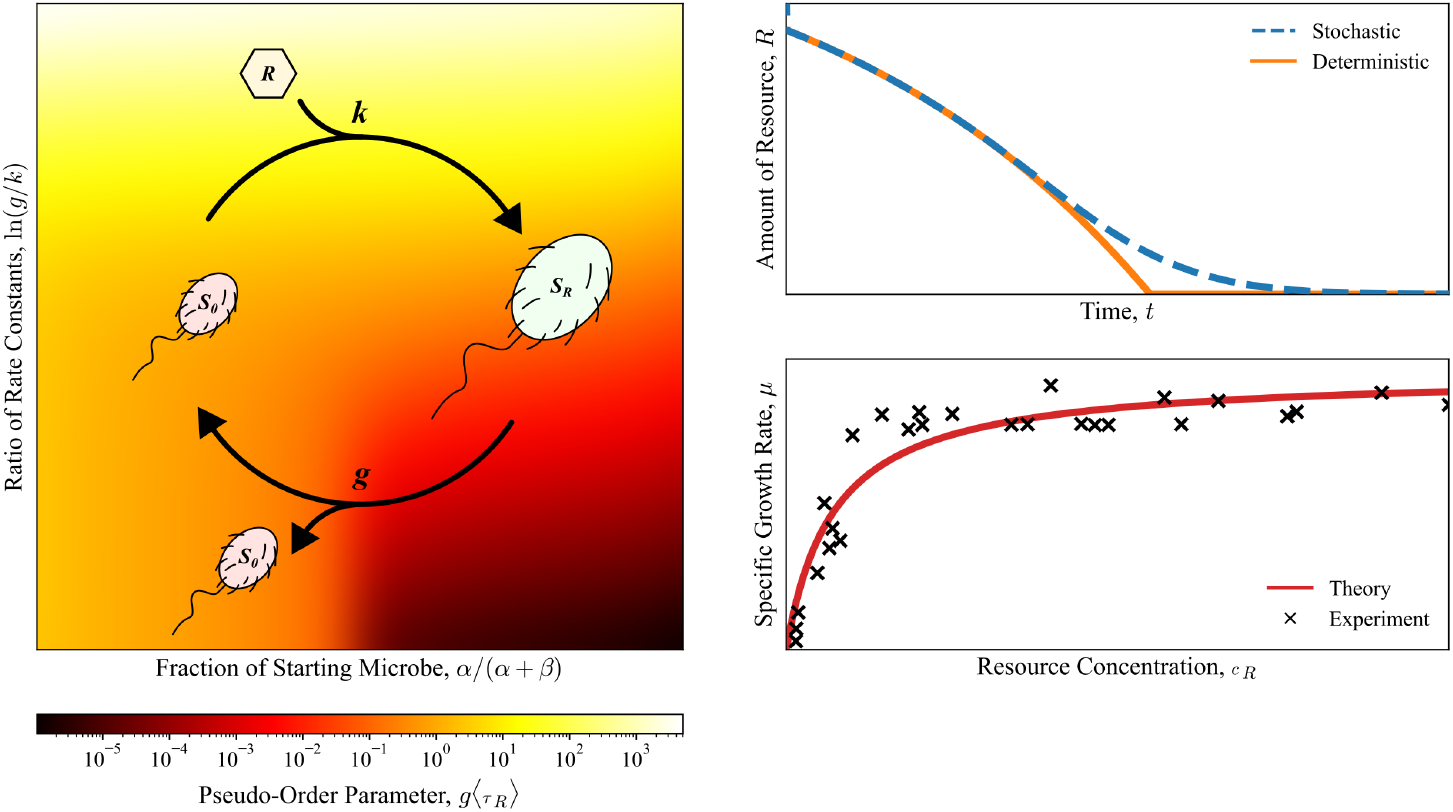

## Introduction

Microbes have the ability to organize into highly elaborate communities as intricate as the ecosystems of higher organisms despite their relatively simple physiology [1, 2]. It is widely believed that this is the result of a large number of microscopic biochemical processes and mesoscopic microbial interactions such as resource competition, predation, and cross-feeding [3–9]. From these multiscale interactions, complex collective properties emerge, which ultimately govern the spatial assembly, behavioral responses, and ecological structures of the resulting consortia [10–13]. In recent years, microbial systems have been intensively studied using a variety of experimental and theoretical techniques, leading to strong advances in our understanding of their dynamics and collective behavior [2, 11, 14–19]. Nevertheless, many aspects of microbial dynamics, like key driving forces that underpin them, remain poorly understood [1, 20, 21].

To support their survival and growth, microbes have to acquire essential nutrients and metabolites from their surroundings. Although many models offer an empirical connection between resource availability and the growth rate of microbes, they often lack clear mechanistic foundations [22, 23]. So, the search for a comprehensive theory that couples resource consumption by microbes to their growth dynamics is still ongoing. Interestingly, complex microbial communities frequently display surprising levels of universality [12, 24]. One example is the apparent hyperbolicity of their growth curves with respect to resource availability — a feature shared by vastly different microbial communities across highly variable chemical and environmental conditions [25, 26]. Such observations suggest the existence of some common, conserved mechanisms that control microbial dynamics, and exploring them can reveal the essential physics of microbial collective behavior [27–29].

Significant theoretical efforts have been devoted to analyze the complex dynamic behaviors of microbial systems [11, 15]. These approaches can usually be divided into three classes: phenomenological kinetic models, deterministic differential equations models, and metabolic flux models [1, 14, 15]. The first kind encompasses growth laws derived to fit experimental measurements, which borrow ideas from kinetic analysis of chemical reactions. Models belonging to this category include the ones famously studied by Monod [25], Droop [30, 31], and many others [23]. While this approach often can capture real-world observations reasonably well, it remains a black box by construction, and thus offers limited mechanistic insights. Besides, there are situations wherein these methods exhibit considerable inconsistencies [32]. The second kind maps ecological interactions onto a set of coupled, deterministic differential equations tracking population dynamics [33]. The inherent continuity of their solutions unfortunately means that they cannot properly describe extinction events [14, 34], at least without resorting to more mathematically involved formulations [35– 37]. The third kind builds on metabolic reaction networks to compute steady-state flux distributions [15]. Flux balance analysis serves as one key example, where an objective function pertaining to some biological functions of interest is optimized subject to mass-balance and capacity constraints [38–40]. Such treatments are desirable in systems microbial ecology, but their huge parameter sets risk over-parameterization [41], and the method’s overall behavior may turn out to be insensitive to the exact values of these parameters [42].

Moreover, most of these theoretical approaches neglect an important property of microscopic systems: stochasticity. When processes occur in a system consisting of only a small number of participating particles, random fluctuations from their collisions become so significant that their behaviors might deviate from the ensemble average, particularly in dissipative systems like cellular environments [43–45]. Some of these noises amplify as they propagate along biochemical pathways, eventually becoming strong enough to influence the outcomes of biological processes [46, 47]. It is therefore not a surprise that, granted sufficiently fine experimental measurements can be made, all cellular activities should manifest in a probabilistic manner [48]. These include the precise mechanisms governing how cellular resources are transported to microbes [43, 49, 50], and how cells grow and divide [51–54]. Although attempts have been made to connect stochastic cellular processes to population-level dynamics, they remain either too complex to be analytically tractable [54], or too phenomenological [55], leaving the key drivers of microbial community dynamics largely unelucidated.

Here, we propose a stochastic model of microbial ecological dynamics under minimal assumptions to explain how mesoscopically deterministic observations of microbial dynamics emerge from microscopic processes. In our approach, the complex metabolic network of microbial cells is mapped onto an effective two-step mechanism: a resource absorption step, followed by cell division. Using analytical and numerical calculations supported by extensive Monte Carlo simulations, we develop a general theory of single-species-single-resource microbial growth dynamics, evaluate the effects of stochasticity, and reproduce experimental observations and well-known empirical relations like the Monod equation. Our results show that, with as few as two constants specifying the rates of resource consumption and cell division, a wide spectrum of collective behaviors in microbial populations can be faithfully recapitulated and explained using simple physical-chemical arguments.

## Model and Methods

To understand the coupling between resource consumption by microbes and their population growth, we propose a minimal theoretical model as presented in Fig. 1A. In this approach, microbial dynamics is mapped onto a two-step reaction involving three types of particles: the unfed microbes labeled as *S*_0_, the resource molecules labeled as *R*, and the fed microbes labeled as *S*_*R*_. It is assumed that only the fed microbes *S*_*R*_, which have already absorbed a resource particle, can divide to increase the microbial population. The overall kinetic scheme for this process can be written as

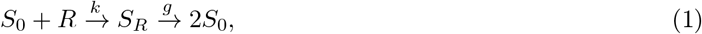

where *k* and *g* are the feeding and the growth rate constants, respectively. The rate constants *k* and *g* are coarse-grained parameters that combine multiple biochemical processes relevant to the consumption and division stages. It is natural to first study this model in the simplified setting where only one type of microbial species feeds on one type of limiting resource, as having a thorough understanding of this system would allow us to extend it to the more realistic multiple-species-multiple-resource case with additional ecological interactions, or to account for more biochemical details.

**Figure 1:**
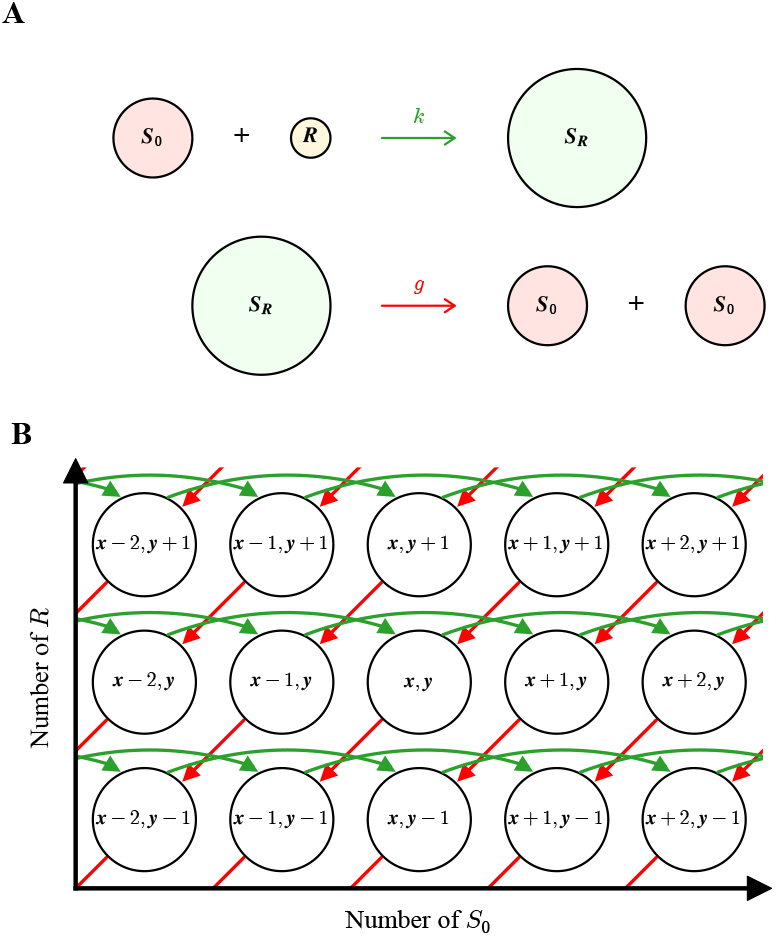
Overview of the minimal stochastic model of microbial ecological dynamics in the single-species-single-resource setting. (**A**) Schematic view of the stochastic model for microbial resource consumption and growth via cell division. The consumption rate constant is *k*, and the division rate constant is *g*. (**B**) Microbial dynamics as a set of stochastic transitions between discrete states. The state (*x, y*) corresponds to the case wherein the system contains *x* microbes of type *S*_0_ and *y* resource molecules *R*. Red arrows correspond to resource consumption with the rate constant *k*, and green arrows correspond to microbe growth with the rate constant *g*.

We denote 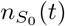, *n*_*R*_(*t*), and 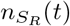 as the number of particles of type *S*_0_, *R* and *S*_*R*_ at time *t*, respectively. Initially, at *t* = 0, it is assumed that there are *α* microbes of type *S*_0_, *β* resource molecules *R*, and no microbes of type *S*_*R*_, setting the initial conditions of the system as

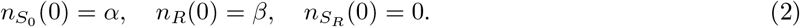

The stoichiometric relations outlined in Eq. (1) and the initial conditions specified in Eq. (2) entail the conservation law

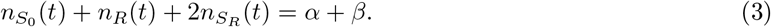

Eq. (3) indicates that there are only two independent degrees of freedom in our system, and at any moment the state of the system can be fully specified by knowing the amounts of any two of the three components (Fig. 1B). We study this model using analytical and numerical calculations supported by extensive Monte Carlo computer simulations. All mathematical details involved are presented in the Supplementary Information.

## Results and Discussion

### General Stochastic Description of Microbial Ecological Dynamics

#### Chemical Master Equation

To simplify the notations, we define 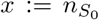, *y* := *n*_*R*_, and 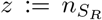, and we introduce *P*_*x,y*_(*t*) as the probability of finding the system in the state (*x, y*) with *x* microbes of type *S*_0_ and *y* resource molecules *R* at time *t*. The amount of fed microbes *S*_*R*_, *z*, can always be found from Eq. (3) by

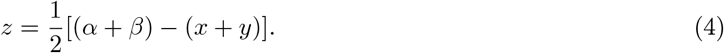

Hence, time evolution of the composition of our reaction system can be viewed as a set of stochastic transitions between discrete states on a two-dimensional lattice, such that its dynamics is governed by the chemical master equation

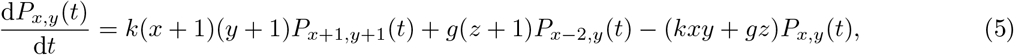

as outlined in Fig. 1B. In Eq. (5), the first term corresponds to the uptake of resource particles *R* by microbes of type *S*_0_ with the rate constant *k*, and the second term corresponds to cell division of the microbes of type *S*_*R*_ with the rate constant *g*. Both of these contribute to increasing the probability for the system to stay in state (*x, y*), while the third term lowers it by accounting for the transitions out of (*x, y*).

Since Eq. (5) is generally not analytically solvable due to its nonlinearity [56], a stochastic simulation algorithm is implemented to study the system’s evolution in a numerically exact way [57]. Details of the computational procedure, as well as the relevant mathematical background of our model, can be found in Section S1 of the Supplementary Information. Fortunately, approximate closed-form analytical solutions of Eq. (5) can be obtained if timescale separation between the pair of coupled reactions is permissible. This happens when one of the rate constants is exceedingly larger than the other, and we call the distinct conditions under which we can reduce Eq. (5) into a set of exactly solvable effective master equations limiting regimes.

#### Analytical Solutions to the Effective Master Equations in Limiting Regimes

There exist three limiting regimes wherein Eq. (5) can be approximated analytically, allowing us to fully analyze microbial ecological dynamics through the lens of our proposed microscopic mechanism. The first is the fast-growing regime when *g* ≫ *k* for any initial conditions, the second is the resource-limited fast-feeding regime when *g* ≪ *k* for *α* ≥ *β*, and the third is the species-limited fast-feeding regime when *g* ≪ *k* for *α < β*. In each of these situations, the microbial dynamics of the system can be described by a set of effective one-dimensional stochastic processes, as illustrated in Fig. 2 and Fig. S2.1. Mathematical validity of these projections is discussed in Section S2 of the Supplementary Information. For clarity, the corresponding limiting regimes and the quantities associated with them are denoted by superscripts A, B, and C, respectively. Regimes with multiple dynamic behaviors, as explained below, are further distinguished by Roman numerals.

**Figure 2:**
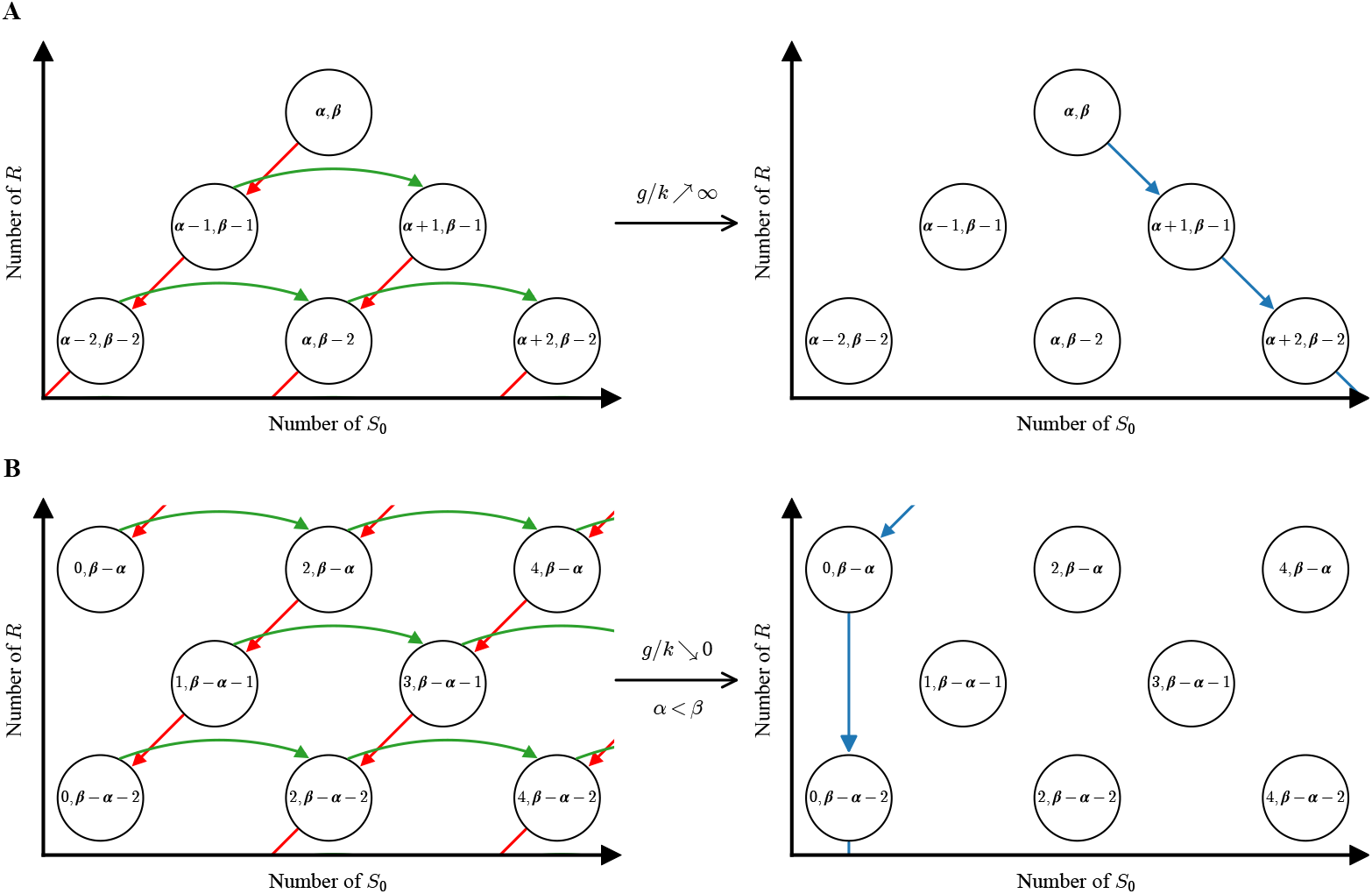
Projections of stochastic microbial dynamics in the limiting regimes into effective one-dimensional transitions. (**A**) Fast-growing regime (*g* ≫ *k*). (**B**) Species-limited fast-feeding regime (*g* ≪ *k, α < β*).

Let us start with the case when the growth rate is very fast (*g* ≫ *k*). In this limiting regime, after each unfed microbe *S*_0_ absorbs a resource particle *R*, it instantaneously disintegrates into two new unfed microbes of type *S*_0_. In other words, *S*_*R*_-type microbes only appear transiently with their amounts being effectively zero. This is equivalent to considering the second-order reaction 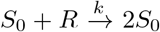, where the production *S*_*R*_ quickly reaches a stationary state at *z* = 0. Such a physical interpretation is similar to the standard quasi-steady-state approximation used in mean-field chemical kinetics. Accordingly, the conservation relation in Eq. (4) reduces to

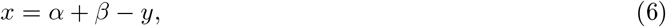

with 0 ≤ *y* ≤ *β*. This means the system can be fully described by a single random variable, say *y*. Fig. 2A shows how the original set of two-dimensional transitions can be mapped onto a set of effective, one-dimensional transitions. For example, if we start from the initial state (*α, β*), then based on Eq. (1) the system can only transitions to the state (*α* − 1, *β* − 1), after which it immediately transitions into the state (*α* + 1, *β* − 1) because *g* ≫ *k*. As a result, these two transitions can be viewed as one effective, virtual transition from the state (*α, β*) to the state (*α* + 1, *β* − 1). This continues until the system runs out of resources *R* (Fig. S2.1).

The effective master equation in this case can be written as

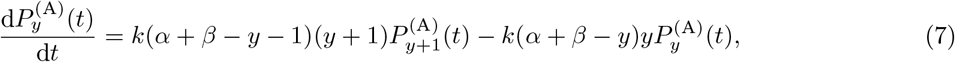

and the solution of Eq. (7) is given by

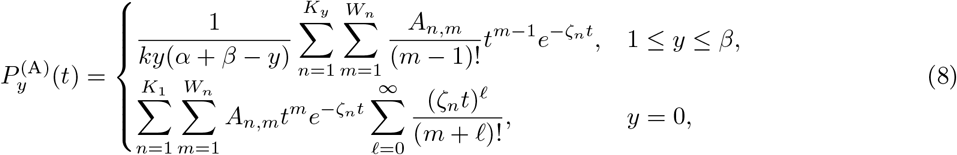

where the coefficients *K*_1_, *K*_*y*_, *W*_*n*_, *A*_*n,m*_ and the factors 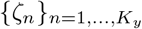 come from generalized partial fraction decomposition in the Laplace-transformed domain [58]. The derivation of Eq. (8) can be found in Section S3 of the Supplementary Information.

One can then easily obtain a full description of microbial dynamics in this limiting regime, as the *n*-th moment of the number of each species can be found from

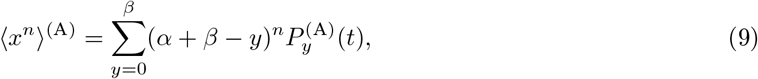

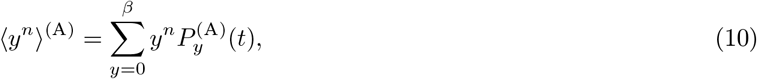

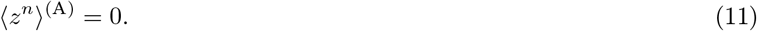

Analytical calculations of the time evolution of mean quantities of microbes and resources in the fast-growing limit are presented in Fig. 3A. As can be seen, the amount of *S*_0_-type microbes quickly increases with time, reaching a plateau when all resource molecules *R* are consumed, while the amount of *S*_*R*_-type microbes effectively stays at zero. It is important to stress that ⟨*z*⟩^(A)^ = 0 does not mean that no *S*_*R*_-type microbes physically exist in this limit. Instead, it means *S*_*R*_-type species only persist on infinitesimally narrow time intervals, such that the average of their quantity is essentially zero.

**Figure 3:**
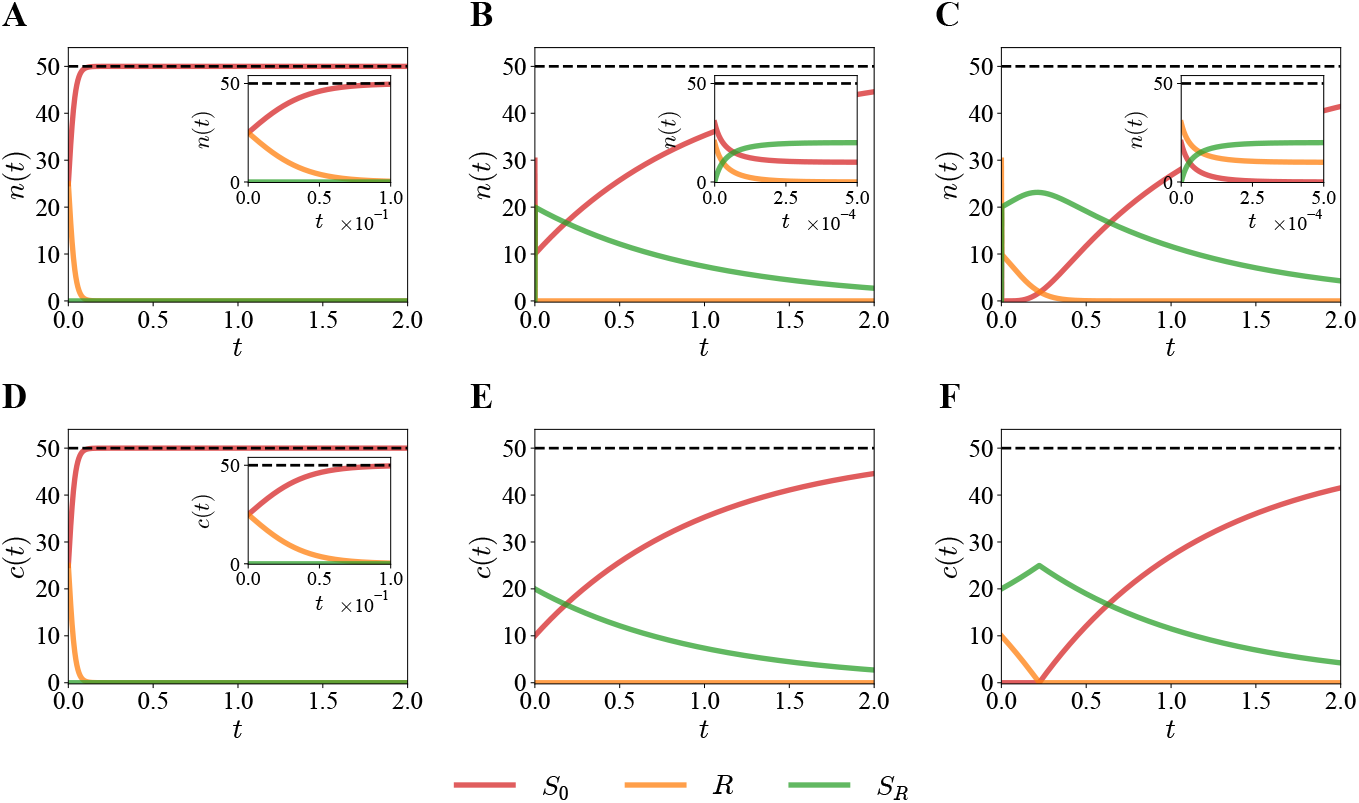
Analytical calculations of microbial dynamics in the limiting regimes. Time evolution of the three key species *S*_0_ (red), *R* (orange), and *S*_*R*_ (green) are presented. The black dashed line marks the conserved quantity *α* + *β*. (**A**–**C**) Mean number of species *n*(*t*) obtained from the stochastic framework. (**A**) Fast-growing limit (*g* ≫ *k*) with parameters *k* = 1 s^−1^, *g* = 10^3^ s^−1^, *α* = *β* = 25. (**B**) Resource-limited fast-feeding limit (*g* ≪ *k, α* ≥ *β*) with parameters *k* = 10^3^ s^−1^, *g* = 1 s^−1^, *α* = 30, *β* = 20. (**C**) Species-limited fast-feeding limit (*g/k* ≪ 1, *α < β*) with parameters *k* = 10^3^ s^−1^, *g* = 1 s^−1^, *α* = 20, *β* = 30. Insets show the early-time dynamics. (**D**–**F**) Concentrations of species *c*(*t*) obtained from the deterministic framework using the same parameters as in (**A**–**C**), respectively. Insets in (**A**–**D**) show expanded views of the initial rapid consumption or generation of molecules.

Another limiting regime is when resource consumption is much faster than cell division (*g* ≪ *k*), but the starting number of resource molecules *R* is smaller than or equal to the starting number of *S*_0_-type microbes (*α* ≥ *β*). It can be argued that the system’s behavior changes abruptly once all resources *R* are purged from the system (Fig. S2.1A). Equivalently, we say that, this system can switch between two distinct dynamic behaviors, and the times which either behavior manifests are separated by some finite critical time *τ*_*R*_. Physically, *τ*_*R*_ is the first instance all resources *R* become extinct.

For *t < τ*, only the process of resource absorption takes place, 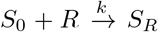, until all resources *R* are consumed. Only one random variable is needed to specify the state of the system because the increase in the amount of *S*_*R*_ is exactly equal to the decrease in the amount of *S*_0_ or *R*. For convenience, we choose to follow the variable *y*, and we have

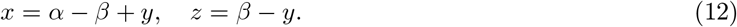

The corresponding effective master equation can be written as

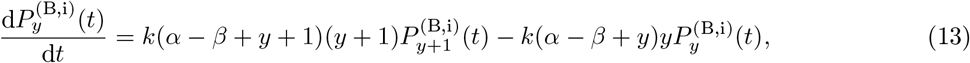

where i labels the dynamic behavior for *t < τ*_*R*_. It can be shown that the solution of Eq. (13) is

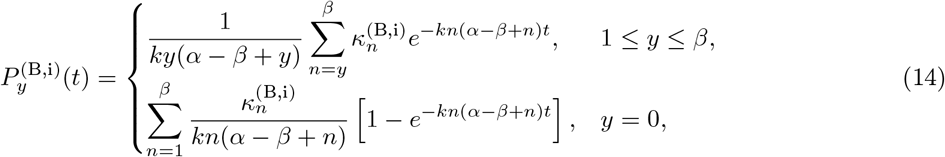

where the coefficients 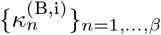 are obtained from the partial fraction decomposition of nondegenerate poles in the Laplace-transformed domain, as derived in Section S4 of the Supplementary Information.

Behavior of the system changes for *t* ≥ *τ*_*R*_ as all resources vanish (*y* = 0), meaning only the cell division process can take place, 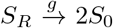. Again, only one random variable is needed to describe the state of the system, and we choose *z* with

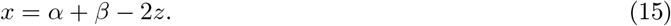

The corresponding effective master equation can be written as

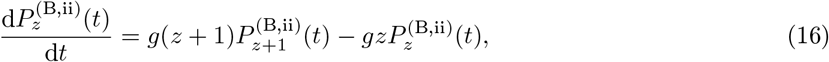

where ii labels the dynamic behavior for *t* ≥ *τ*_*R*_. The solution of Eq. (16) is

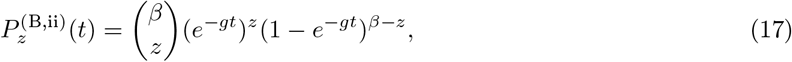

which is a known result for linear pure-death processes [56, 59].

To properly evaluate the microbial dynamics in this limiting regime, it is necessary to account for the fact that *τ*_*R*_ is itself a continuous random variable with a well-defined probability density function 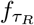 With it, the *n*-th moment of the number of each species can be found from

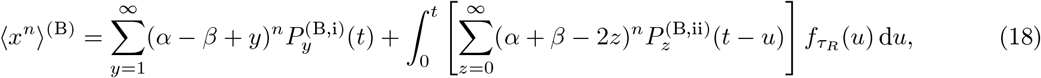

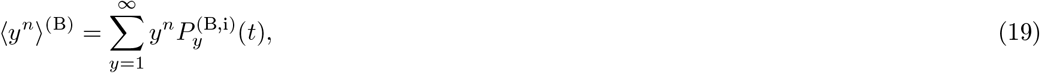

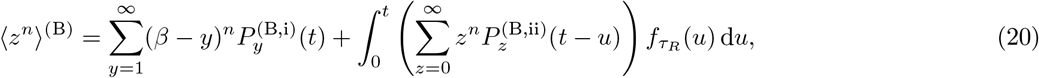

where 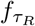 and the integrals in Eqs. (18) and (20) are written explicitly in Section S5 of the Supplementary Information. Fig. 3B shows the time evolution of the mean quantities of each species in this limiting regime. One can see that, at very early times, the amount of *S*_0_-type microbes decreases rapidly, then increases until saturation at later times. The population of *S*_*R*_-type microbes exhibit the opposite dynamics: at very early times, their amount increases rapidly; after all resources are consumed, their number starts dropping.

The third limiting regime is when the consumption of resources is again much faster than cell division (*g* ≪ *k*), but the starting number of resource molecules *R* is strictly larger than the starting number of *S*_0_-type microbes (*α < β*). As shown in Fig. 2B, the effective dynamics in this limit is again one-dimensional, and one can identify three different dynamic behaviors separated by the first extinction times 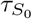 and *τ*_*R*_ from Fig. S2.1B. For 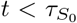, only the resource consumption process takes place until all *S*_0_-type microbes are consumed, 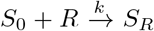. For 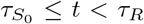, the *S*_*R*_-type microbes start to slowly divide into *S*_0_-type microbes, which, upon creation, immediately transform into *S*_*R*_ by utilizing the remaining resources *R* in the system. This means that at these times, *x* = 0 effectively, and this situation continues until all resources are consumed. It can be described as 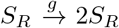. Finally, for *t* ≥ *τ*_*R*_, there are no resources left (*y* = 0), and the reminaing *S*_*R*_-type microbes decay into *S*_0_-type microbes, 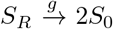. The dynamic behaviors for 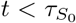 and for *t* ≥ *τ*_*R*_ are analogous to the corresponding cases in the resource-limited fast-feeding regime. As for 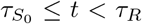, the system behaves in ways different from the previous limiting regimes, but its joint probability distribution can still be written analytically.

Due to the added complexity that the effective state spaces in this limiting regime depends on the parity of the difference *β* − *α*, the effective master equations and their solutions — labeled by i, ii and iii as 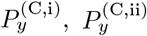, and 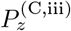, respectively — assume highly nontrivial forms. Thus, we present the mathematical details of their derivations in Section S6 of the Supplementary Information. The *n*-th moment of the number of each species can be found from

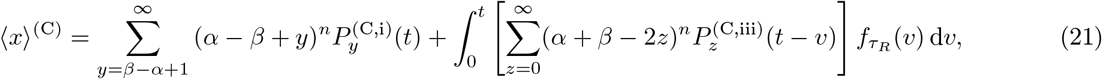

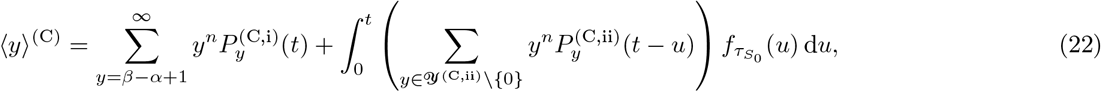

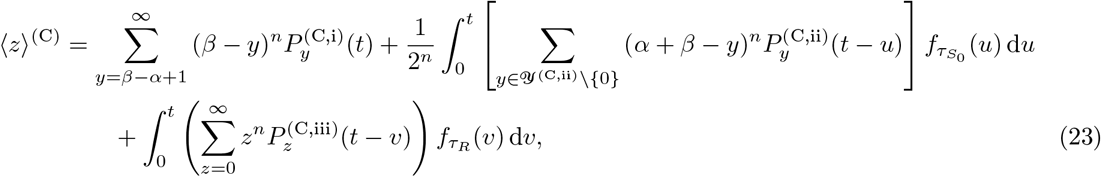

as derived in Section S7 of the Supplementary Information. In particular, *Y*^(C,ii)^ denotes the set of allowed *y*-values in the effective state space of phase ii of regime C. The time evolution of the mean quantities of each species in this limiting regime is shown in Fig. 3C. As outlined above, one can discern how the dynamic behavior of the system is segregated into three phases. For instance, the mean of *S*_0_-type microbes is seen to decrease rapidly at first, then for some time stays close to zero, and finally begins to surge and saturates.

### Deterministic Description of Microbial Ecological Dynamics

When the number of microbes and resource molecules is large, it is more convenient to describe the composition of the system in terms of concentrations, defined by

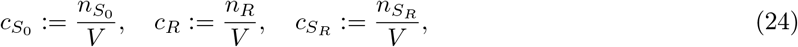

where *V* is the volume of the system. Then, dynamics of the system can be analyzed using the chemicalkinetic-like deterministic differential equations

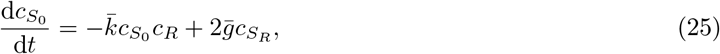

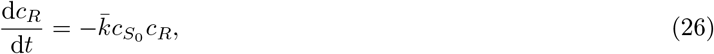

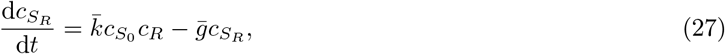

where 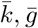 are the macroscopic rate constants. It is also helpful to consider the total concentration of all microbes, 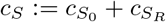, whose time evolution obeys

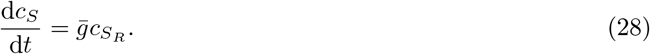

Similar to the stochastic framework, the initial conditions in the large-system-size limit are taken to be

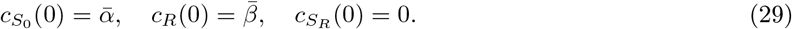

Generally, Eqs. (25)–(27) cannot be solved analytically. However, similar to the stochastic approach, we can analyze the three limiting regimes wherein their exact solutions can be analytically approximated. The derivations are presented in Section S8 of the Supplementary Information.

In the fast-growing limit 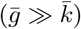, the quasi-steady-state approximation can be applied to get

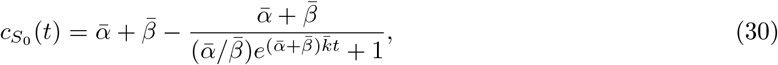

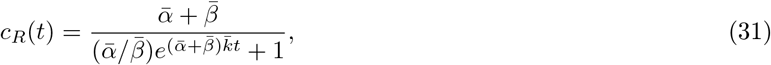

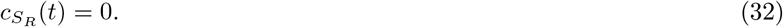

The analytical approximations in this limit are presented in Fig. 3D. One can see that the predicted deterministic dynamics in the fast-growing limit is essentially identical to the results from our more general stochastic analysis, by comparing with Fig. 3A. Thus, in this limiting regime, it is justifiable to use the deterministic approach.

In the resource-limited fast-feeding regime (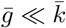 and 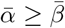), the corresponding analysis yields

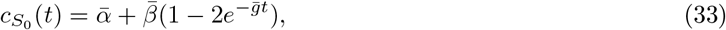

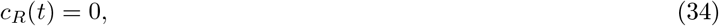

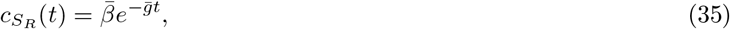

for *t >* 0. The results of our calculations are presented in Fig. 3E. Compared with the stochastic case in this limit (Fig. 3B), the deterministic chemical-kinetic approach describes the dynamics of the system well for most times except near *t* = 0, when the initial decay of the resource molecules is not captured.

Finally, in the species-limited fast-feeding regime (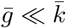 and 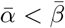), we have

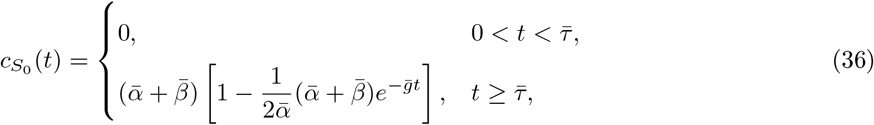

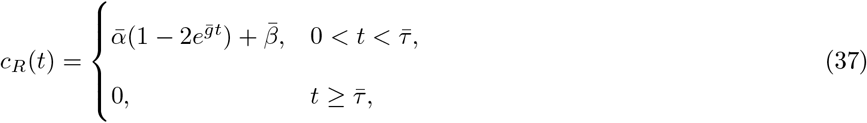

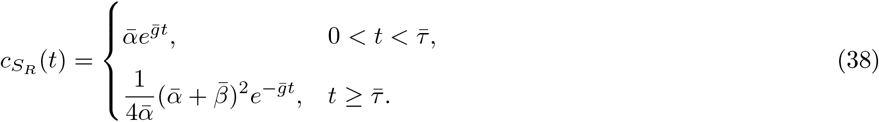

In these expressions, the characteristic time 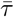 is given by

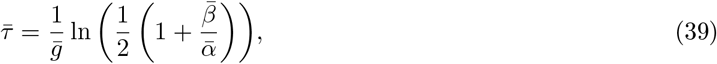

which is the time at which all resources are consumed. It is important to note that these analytical approximations are constructed piecewise to circumvent the atto-fox problem [34]. The actual numerical solutions of Eqs. (25)–(28) should in fact be differentiable everywhere on its domain.

The results of our calculations in this limit are presented in Fig. 3F. Comparing with more general stochastic approach in Fig. 3C, we notice that although the deterministic method qualitatively agrees with the stochastic results in most situations, there are two major differences. The chemical-kinetic-like calculations miss the early-time dynamics by construction, similar to the fast-feeding resource-limited case, and predict a much sharper transition at 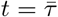.

### Dynamic Phase Diagram of Microbial Ecological Dynamics

Our stochastic approach enables us to better understand how the intricate interplay between resource consumption by microbes and their divisions drives microbial ecological dynamics in the single-species-single-resource setting. Since Eq. (5) cannot be solved analytically for general conditions, we use computer simulations to investigate the dynamics of the system numerically exactly. Together with our analytical results discussed previously, we are able to comprehensively study the distinct behavioral shifts exhibited by the system at finite times.

Quantitatively, we choose two quantities to specify the system’s dynamic transitions. The first quantity is the mean first extinction time of resource molecules, ⟨*τ*_*R*_⟩. In the limiting regimes, these times can be written as

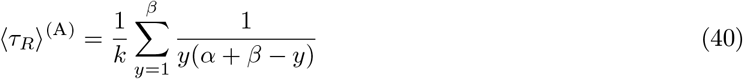

for the fast-growing regime (*g* ≫ *k*),

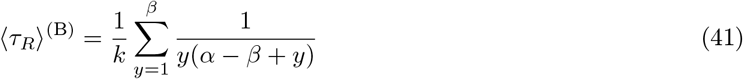

for the resource-limited fast-feeding regime (*g* ≪ *k* and *α* ≥ *β*), and

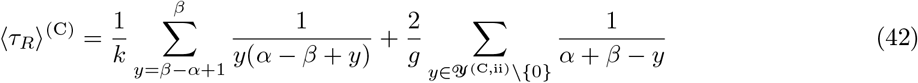

for the species-limited fast-feeding regime (*g* ≪ *k* and *α < β*). But in general, these times can only be sampled from computer simulations. The second quantity is the fractions of each particle type, defined by

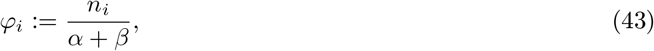

for *i* ∈ {*S*_0_, *R, S*_*R*_}. In addition, our previous arguments suggest that microbial dynamics strongly depends on the relative magnitudes of the feeding and growth rate constants. For this reason, we interpret the ratio *g/k* as the effective field that controls the behavior of the system.

Fig. 4 illustrates the results of our theoretical analysis of microbial dynamics over a wide range of parameters, which combines explicit analytical calculations in the limiting regimes with extensive Monte Carlo computer simulations in the intermediate cases. We find that relative magnitude of the feeding and growing rate constants has a strong effect on the dynamics of resource consumption. Starting from Fig. 4A, which encompasses the *α* ≥ *β* case, we see that the mean first extinction time of *R* stays almost constant when *g < k*. But as soon as *g* becomes larger than *k*, an abrupt crossover occurs, with ⟨*τ*_*R*_⟩ quickly decreasing until reaching another constant value. The dynamics is slightly different in the *α < β* case (Fig. 4B), where ⟨*τ*_*R*_⟩ decreases with increasing *g/k* until reaching a small constant value for *g > k*. Still, deviation between the analytical calculations and simulated results exists near *g* = *k*, as highlighted in the inset of Fig. 4B.

**Figure 4:**
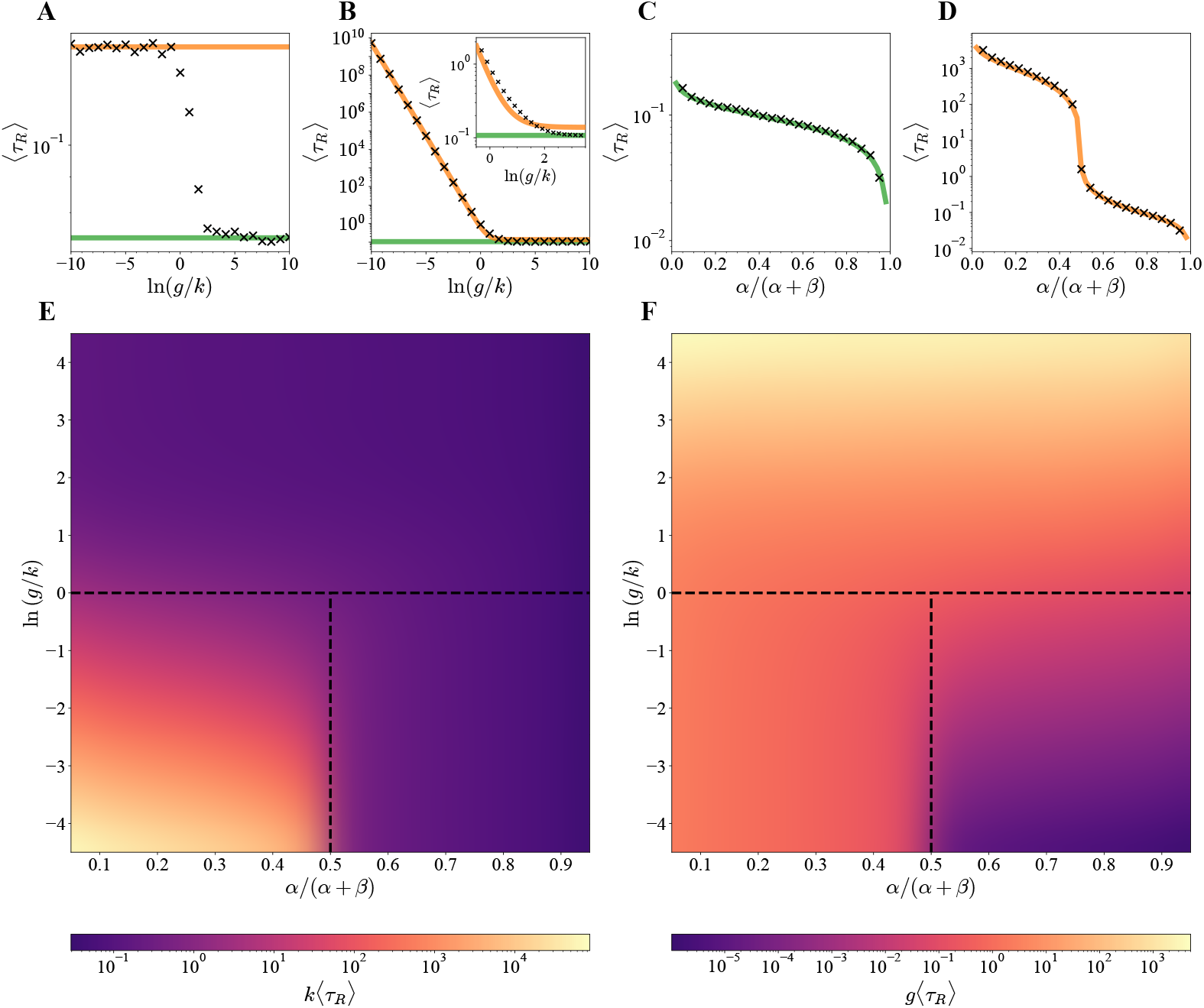
Dynamics of the minimal stochastic model over the parameter space. (**A**) Mean first extinction times of resource ⟨*τ*_*R*_⟩ as a function of the ratio *g/k* for *α* = 35 and *β* = 15 (*α* ≥ *β*). (**B**) Mean first extinction times of resource ⟨*τ*_*R*_⟩ as a function of the ratio *g/k* for *α* = 15 and *β* = 35 (*α < β*). (**C**) Mean first extinction times of resource ⟨*τ*_*R*_⟩ as a function of the initial fraction of *S*_0_ for *g/k* = 10^3^. (**D**) Mean first extinction times of resource ⟨*τ*_*R*_⟩ as a function of the initial fraction of *S*_0_ for *g/k* = 10^−3^. (**E**) Phase diagram of microbial dynamics expressed as in the dimensionless quantity *k*⟨*τ*_*R*_⟩ over the parameter space spanned by the ratio *g/k* and the initial fraction of *S*_0_. For calculations, *k* and *α* + *β* are fixed at 1 and 50, respectively. (**F**) Phase diagram of microbial dynamics expressed as dimensionless quantity *g*⟨*τ*_*R*_⟩ over the parameter space spanned by the ratio *g/k* and the initial fraction of *S*_0_. For calculations, *g* and *α* + *β* are fixed at 1 and 50, respectively. In (**A**–**D**), symbols indicate results from Monte Carlo computer simulations, and orange and green lines correspond to analytical results in the fast-feeding and the fast-growing limits, respectively.

A complementary perspective is to look at how the mean first extinction time of *R* varies with the initial fraction of *S*_0_-type microbes when we fix the ratio *g/k*. These results are presented in Fig. 4C for *g > k* and Fig. 4D for *g < k*. In both cases, ⟨*τ*_*R*_⟩ decreases as we increase the initial fractions of *S*_0_ species since, initially, there are less available resource molecules *R* in the system. However, the first derivative of ⟨*τ*_*R*_⟩ depends strongly on the ratio *g/k*. For *g > k* (Fig. 4C), the rate of decrease holds a steady value; for *g < k* (Fig. 4D), the rate of decrease exhibits a sudden jump at *α* = *β*, making ⟨*τ*_*R*_⟩ plummets rapidly.

These observations allow us to construct effective phase diagrams for the dynamics of our single-species-single-resource model, as presented in Figs. 4E and 4F. Three dynamical regimes can be identified as quantified by the mean first extinction times ⟨*τ*_*R*_⟩. Our theoretical approach can explain these phase diagrams. To be more specific, consider Fig. 4E. When *g > k*, cells divide faster than they can take in resources. This means, soon after the first batch of *S*_*R*_-type microbes is produced, they quickly split into new *S*_0_-type microbes, thereby accelerating the consumption of resources. As a result, the mean first extinction times ⟨*τ*_*R*_⟩ generally remain small. This is the dynamic phase that occupies the upper half-plane in the phase diagram. Two distinct situations are observed in the lower half-plane. When *g < k*, cells absorb resources quicker than they can divide. So, in a relatively resource-poor environment (*α* ≥ *β*), all available nutrients are quickly consumed, such that the mean first extinction times ⟨*τ*_*R*_⟩ stay small. This is the dynamic phase that occupies the lower right quarter of the phase diagram. Majority of biological systems are probably found in this dynamic phase, as we argue later. On the other hand, in a relatively resource-rich environment (*α < β*), a small portion of the initial *S*_0_-type microbes transform into *S*_*R*_-type microbes quickly, but then the reaction system stagnates because regeneration of *S*_0_-type microbes proceeds slowly. This means it takes significantly longer time for the system to run out of resources. As a result, the mean first extinction times ⟨*τ*_*R*_⟩ are large. This is the dynamic phase that occupies the lower left quarter of the phase diagram.

Naturally, these three dynamical regimes can be viewed as extensions of the three limiting regimes under more relaxed values of rate constants *k* and *g*. In fact, from Fig. S9.1, one can see that, generally, our limiting analytical approximations are qualitatively accurate descriptors of microbial dynamics over a wide range of rate constants *k* and *g*, except for a narrow sliver of the phase space where abrupt transitions occur.

### Comparison between the Stochastic and Deterministic Descriptions of Microbial Ecological Dynamics

Microbial dynamics is frequently analyzed by mean-field approaches that use deterministic differential equations to model the time evolution of the concentrations of microbial species and resource molecules in a reaction vessel. However, given the stochastic nature of how microscopic processes operate, it remains an open question as to whether these approaches are mechanistically realistic. Our theoretical model allows us to quantitatively investigate this. We evaluate the changes in the concentrations of microbial species and resource molecules using Monte Carlo computer simulations and compare them with the predictions made from numerically exact solutions of the mean-field chemical-kinetic equations.

From Fig. S9.2A, we see that stochastic and deterministic descriptions of microbial dynamics generally coincide over the whole parameter space, but there are still obvious discrepancies between the framework when *g < k*. The results of our analysis are presented in Fig. 5, where large deviations are observed when the system changes between different dynamic behaviors. In particular, while the deterministic approach predicts sharp transitions, the stochastic approach predicts smoother transitions. Though, as expected, increasing the number of species *α* + *β* in the system improves the agreement between stochastic and deterministic calculations [60–62]. This is quantitatively captured in Fig. S9.2B, which shows that deviations between the stochastic and deterministic curves decreases with increasing system size in a power-law fashion.

**Figure 5:**
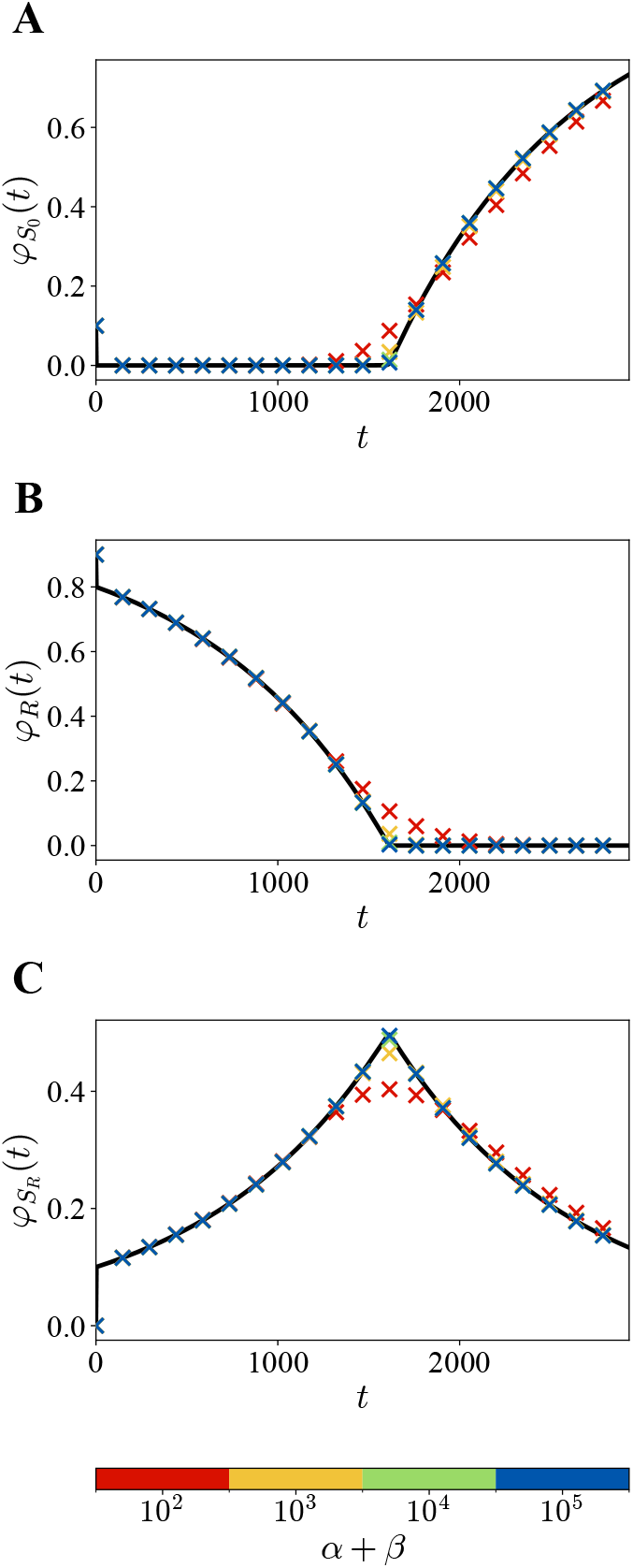
Comparison of temporal evolution of population of species using stochastic and deterministic approaches. (**A**) Fraction of *S*_0_ microbes, 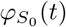. (**B**) Fraction of resource molecules *R, φ*_*R*_(*t*). (**C**) Fraction of *S*_*R*_ microbes, 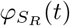. Solid black curves correspond to the exact numerical solutions to the set of deterministic, chemical-kinetic-like differential equations. Color symbols correspond to results of stochastic simulations with different initial conditions. Results are simulated with parameters *k* = 1 s^−1^, *g* = 10^−3^ s^−1^, and *α/*(*α* + *β*) = 0.1.

These observations suggest that the blind use of deterministic methods might incorrectly describe some aspects of microbial dynamics. For instance, it is a well-known hurdle in population ecology that important ecological events, like extinctions, cannot be realized by solutions of typical differential equation models [34]. Another conclusion is that measurements on the behavioral shifts in a microbial system might provide additional information on the system itself, where sharper transitions indicate a larger number of participating species, while smoother transitions indicate a limiting number of microbes or resources. While experiments are often conducted in the large system-size-limit, the latter interpretation might be useful when studying the microbial ecology of marine planktonic bacteria living in the highly dilute oligotrophic ocean [63, 64], or the behavior of a starting colony of isolated species subjected to ecological succession or serial dilution experiments [65–67].

### Comparison with Experiments

Our theoretical method lumps together a large number of biochemical and biophysical cellular processes into two effective steps. This raises the important question of whether our minimal model suffices to capture the dynamics of real microbial systems. To test this, we fit experimental data on glucose consumption by bacterial species using our analytical approximations to the numerical solutions of mean-field differential equations. In here, we specifically use the deterministic approach because, in these experiments, large amounts of bacterial species and resources have been used.

The results of our analysis are presented in Fig. 6 for three different experiments [68, 69]. It should be noted that our minimal model considers *S*_*R*_ as a complex formed by combining equal parts of *S*_0_ and *R*. This may appear inconsistent with reality, where, for instance, the amount of glucose required to support bacterial replication is on the order of 10^9^ molecules per cell [70]. However, only the uptake of the first few sugar molecules is rate-limiting; subsequent transport proceeds much faster because expression of sugar transporter genes is typically induced by their corresponding agonists [71–73]. Consequently, the feeding reaction in our minimal model implicitly represents this initial second-order rate-limiting step, rendering our setup fundamentally different from the fractional parameter defined in a similar study [74]. Thus, we can directly compare the experimentally observed initial resource concentrations with the fitted 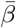 in units of millimolar, as shown in the insets of Figure 6. In all cases, excellent agreement between theoretical predictions and experimental data is observed.

**Figure 6:**
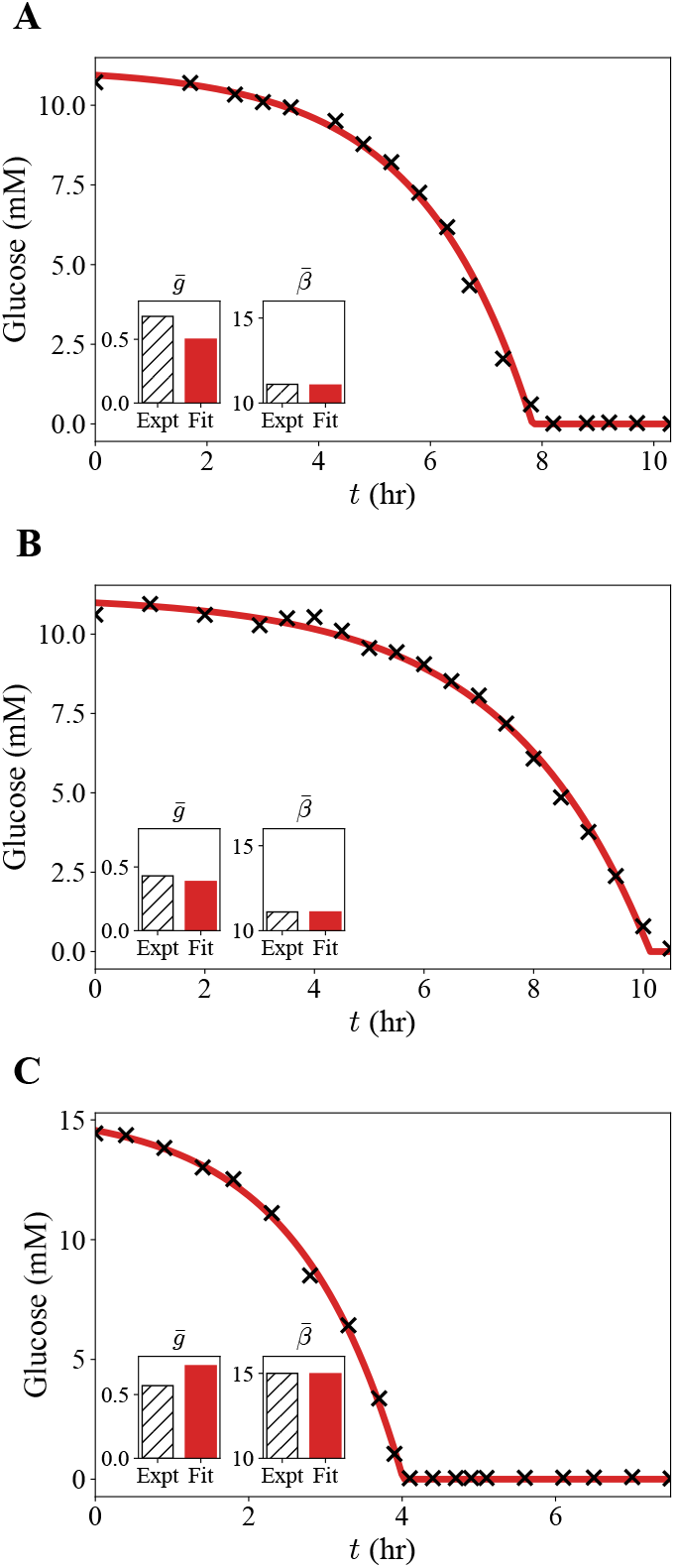
Analysis of glucose consumption in bacterial batch cultures. (**A**) Resource consumption under aerobic conditions as reported by Varma and Palsson [68]. (**B**) Resource consumption under anaerobic conditions by Varma and Palsson [68]. (**C**) Resource consumption in minimal M9 medium by Enjalbert and colleagues [69]. Each red curve corresponds to the best-fit prediction from our model 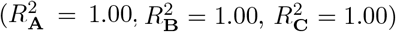. Black symbols correspond to the experimental measurements. The insets compare experimentally measured values of the rate constant 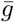 (in units of mM^−1^ s^−1^) and the initial concentration of glucose 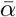 (in units of mM) with the fitted values, with the initial concentrations of bacteria 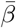 omitted for brevity.

Our fitting of experimental data suggests that the consumption of glucose takes place in the species-limited fast-feeding limit, which is consistent with the procedures specified in the original texts [68, 69]. This means that the resource uptake step is fast while the cell division step is slow, and at the start of all these experiments, the concentration of limiting substrates is greater than the concentration of *S*_0_-type microbes. The fact that our minimal model can successfully describe different experiments on resource consumption by bacterial species indicates that the presented theoretical method can account for relevant universal drivers of microbial dynamics.

### Emergence of Monod Growth Kinetics

One of the most successful and widely utilized approaches to describe microbial dynamics is the phenomenological method proposed by Monod more than seven decades ago [25]. From his empirical observations, he relates the specific growth rate of microbes, defined as

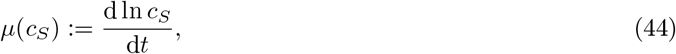

to the total concentration of resources via the equation

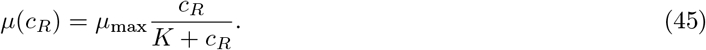

In this expression, the parameter *K* is called the Monod constant, which can be considered as the concentration of resources required for the specific growth rate to reach one-half of the maximal growth rate *µ*_max_.

To show that our minimal theoretical model also captures the hyperbolic growth curve characteristic of Monod kinetics, we explicitly calculate the specific growth rates in the fast-growing and fast-feeding limits, leading to

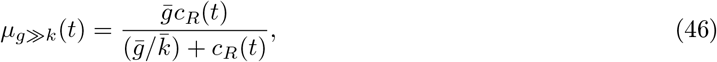

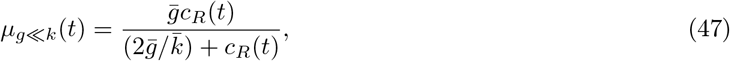

respectively, with derivations provided in Section S10 of the Supplementary Information. At very early times, the values of these growth rates effectively depend on the initial concentration of *R, c*_*R*_(0^+^). Figure 7A shows that both growth curves increase monotonically with *c*_*R*_(0^+^) and saturate asymptotically at 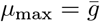. Although the curves approach the same maximum rate and share the same qualitative hyperbolic shape, they exhibit different half-saturation constants *K*. This explains why previous studies assuming *g* ≫ *k* still recovered Monod-like kinetics [74], even though such an assumption might not be biologically accurate. Fitting these expressions to Monod’s original data confirms that *k > g* in general [25]. Since two curves have nearly identical mathematical form, we show only the fit using the most biologically realistic curve *µ*_*g*≪*k*_ in Figure 7B.

**Figure 7:**
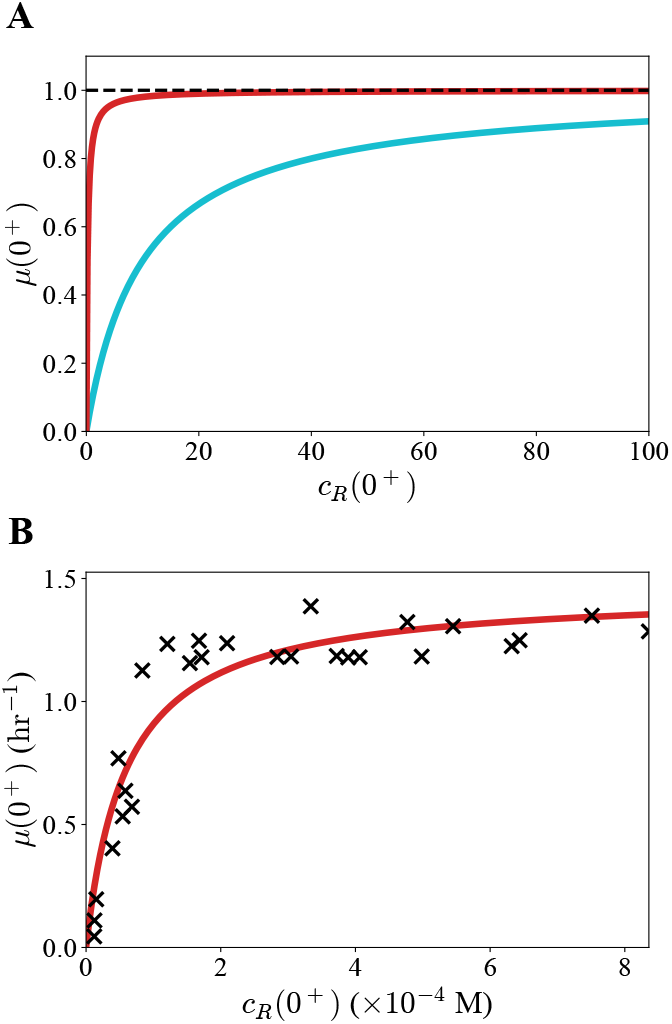
Microbial growth rates as a function of initial concentrations of resources. (**A**) Analytically derived initial growth rate curves in the fast growing (red curve) and fast-feeding (cyan curve) limits. (**B**) Dependence of initial microbial growth rates on initial resource concentrations measured by Monod [25]. Black symbols correspond to the experimental measurements. The fast-feeding curve is used to fit the experimental data (*R*^2^ = 0.89).

As Eq. (45) is a purely phenomenological equation obtained from empirical observations, the physical meaning of the Monod constant *K* has remained hotly debated [22]. Our theoretical approach might offer some possible mechanistic insights into this subject because, using the more accurate description of growth law *µ*_*g*≪*k*_, we obtain

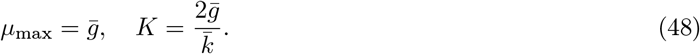

Our theoretical method argues that the maximal rate of microbial growth is limited by the rate of cell division, which is a physically reasonable conclusion [75, 76]. Another theoretical prediction is that the Monod constant *K* actually encodes two important features of the microbial species: the propensity for growth relative to nutrient intake per unit time in the microscopic sense, and the number of offspring produced per division. This aligns with our arguments on why the two-step coarse-graining process is biologically reasonable according to proteome allocation theory in Section S1 of the Supplementary Information [24], and is also conceptually consistent with the recent work by Yamagishi and Hatakeyama showing the universality of hyperbolic growth curves and its possible underlying causes [26].

Accordingly, our rate constants *k* and *g* can be viewed as the coarse-grained outputs of more detailed proteome allocation-based models, which enable us to develop an analytically tractable “effective theory” of microbial dynamics that captures its universal drivers and features. This explains why experimentally measured Monod constants *K* exhibit large variability even between microbial species within the same strain [32]. As the bacterial proteome reorganizes in response to environmental changes [77, 78], the coarse-grained effective rate constants *k* and *g* that represent the outcome of the underlying metabolic networks should also vary significantly, leading to strong variations in *K*. Several computational studies have explored these observations in more detailed frameworks [79, 80], but to the best of our knowledge, our work is the first to provide a general analytical perspective on the problem using a minimal transferable model.

## Conclusions

In this work, we present a minimal stochastic model that enables a full analytical and numerical exploration of single-species-single-resource microbial ecological dynamics. The model reveals three distinct dynamical regimes controlled by the relative magnitudes of the rate constants of feeding and growth, and by the initial conditions. It allows us to investigate how stochastic fluctuations affect microbial dynamics, which become relevant when species or resource numbers are small. Despite its simplicity, the framework not only recovers experimental observations and Monod-like growth kinetic, but also provides a mechanistic interpretation of the maximal growth rate and the Monod constant. This supports the view that universal drivers and features of microbial growth can be understood from a few parameters. In contrast to earlier deterministic treatments that implemented a similar two-step picture but either relied on biologically unrealistic limits [74], or only focused on microbial dynamics under specific conditions [81], our analysis captures the system’s general behavior and identifies the conditions under which deterministic equations fail. Natural extensions of our model include the introduction of multiple species, multiple resources, cross-feeding, or competitive interactions, but the present work suffices to offer an analytically tractable baseline for coupling theory with experiment and for interpreting the microscopic origins of collective microbial dynamics.

## Acknowledgments

A.B.K. was supported by the Welch Foundation (C-1559), the NIH (R01GM148537), and the Center for Theoretical Biological Physics sponsored by the NSF (PHY-2019745). The authors are grateful to B. Ø. Palsson for helpful communications.

## Data Availability

All code files used to perform the analytical calculation, numerical simulation, and figure generation in this study are publicly available at https://github.com/cfaleung1-rice/publication_1r1s under the MIT license. The same repository can be found at https://doi.org/10.5281/zenodo.21252284 [82].

## Supporting information

The following file is available to support our investigation.

- Implementation of the stochastic simulation algorithm, mathematical proofs of timescale separation, derivations of the solutions to the effective master equations and the analytical approximations to the mean-field differential equations, and extra supporting figures.

