## Supplementary Information for "A Minimal Stochastic Model of Microbial Ecological Dynamics in a Single-Species-Single-Resource Setting"

### S1 Mathematical Review and Details of the Stochastic Simulation Algorithm

In this study, we map the complex underlying mechanisms of microbial ecological dynamics onto a two-step chemical-kinetic model with the aim to understand how apparent universality of their behaviors emerges. The fact that vastly different microbial communities exhibit many conserved macroscopic behaviors signifies their invariance to the finer details of each distinct system [1–4]. It is therefore reasonable to consider a transferable minimal model capturing the essence of their mechanistic similarities. Whilst the organizations of microbial ecosystems are inherently complex and high-dimensional, both experimental and theoretical studies have shown that a basic consumer-resource-type model, encoding no direct interactions between individual entities except for exploitation competition on limiting resources, is often sufficient to recapitulate their common emergent properties [3, 5, 6]. Inspired by this, our model focuses on microbial ecological dynamics driven by the coupling between the resource uptake process and the growth and division process, which we coin simply "feeding" and "growth" respectively. In particular, we start from the single-species-single-resource case, from which the generalized many-body case can be reconstituted.

Consider the following key notations:

- $S_0$ – unfed microbe.
- $R$ – resource.
- $S_R$ – fed microbe.
- $k$ – microscopic feeding rate constant.
- $g$ – microscopic growth rate constant.
- $X_t$ – count of $S_0$ at time $t$.
- $Y_t$ – count of $R$ at time $t$.
- $Z_t$ – count of $S_R$ at time $t$.
- $n_i(t)$ – count of particle $i$ at time $t$.
- $\alpha$ – initial count of $S_0$.
- $\beta$ – initial count of $R$.
- $c_i(t)$ – concentration of particle $i$ at time $t$.
- $\tau_i$ – first extinction time of particle $i$.
- $\varphi_i(t)$ – fraction of $n_i(t)$ (resp. $c_i(t)$) normalized by $\alpha + \beta$ (resp. $\bar{\alpha} + \bar{\beta}$).

Also, recall the mechanism

$$S_0 + R \xrightarrow{k} S_R \xrightarrow{g} 2S_0, \tag{S1}$$

and the conservation law

$$X_t + Y_t + 2Z_t = \alpha + \beta. \tag{S2}$$

Please refer to the Main Text for the full set of notations used. Macroscopic version of each term, or in the above case the concentrations $\bar{\alpha} := c_{S_0}(0), \bar{\beta} := c_R(0)$, is indicated by a bar. Also, note that, the symbol $\langle\cdot\rangle$ is used to denote the average of a quantity by conventions of the physical sciences, but $\mathbb{E}[\cdot]$ would be used in the Supplementary Information for mathematical preciseness.

It is important to note that the two-step coarse-graining of a general metabolic network is not merely a convenient mathematical trick for coupling resource flow to population dynamics. Microbial proteomes have in fact been seen to segregate into functionally distinct partitions with a clear supply-demand architecture, and it has been shown that a minimum of three proteomic sectors is enough to recover the linear growth laws relating cell proliferation rates to gene expression levels [7]. Whilst these growth laws are phenomenological in nature [2], several mechanistic studies exist corroborating such findings. For instance, condition-dependent profiling of bacterial proteomes has demonstrated how the metabolic and biosynthetic enzyme expressions are differentially regulated with changing growth rates [8], with recent computational studies bridging the gap between proteomic and transcriptomic modularisation [9]. On the other hand, theoretical studies have shown how proteome efficiency increases along metabolic pathways of nutrient flow [10, 11]. These justify lumping the complex metabolic network into a single effective resource-acquisition module that supplies precursors to a growth-and-division module, which run at distinctly different rates to optimize microbial growth in fluctuating environments [11].

Consequently, we hypothesize that, from our microscopic description, the feeding rate constant $k$ should be much larger than the growth rate constant $g$ for most microbial species. This follows from the observation that microbial doubling rates are typically limited by downstream biomolecular processes, such as ribosomal synthesis and translation, rather than by upstream processes such as nutrient uptake [10, 12]. Although this asymmetry may appear counterintuitive at first, one must consider that maximizing the functional output of a process does not always produce a proportional gain in fitness [13]. One such example is the wild-type expression level of aminoacyl-tRNA synthetases that teeters on a sharp fitness cliff: whilst its underproduction is strongly deleterious, its overproduction confers little additional selective advantage [14]. Moreover, like how eukaryotic cell cycles are regulated by numerous checkpoints to ensure the production of viable offspring [15], bacterial cell cycles incorporate multiple safety features which lower the effective population growth rate under nonideal conditions [16, 17]. Our hypothesis of $k \gg g$ stands in stark contrast to previous theoretical studies that employ a similar two-step coarse-graining scheme and assume $k \ll g$ with lacking biological justification [18, 19]. Nonetheless, our model encompasses the full range of $k$ and $g$ values, and we show that hyperbolicity in growth curves emerges naturally from both limiting cases.

In our model, we consider the sample space

$$\Omega = \prod_{n=0}^{\infty} ((0, \infty] \times \{\mathrm{F}, \mathrm{G}\}), \tag{S3}$$

such that each $\omega \in \Omega$ is a sequence of tuples of the form

$$\omega = \big((s_n, \xi_n)\big)_{n \in \mathbb{N}_{\geq 0}}, \tag{S4}$$

where $s_n \in (0, \infty]$ is the waiting time between each reaction and $\xi_n \in \{\mathrm{F}, \mathrm{G}\}$ is the label of the reaction channel fired, with F and G indicating feeding and growth respectively. The corresponding $\sigma$-algebra $\mathscr{F}$ is

$$\mathscr{F} = \bigotimes_{n=0}^{\infty} (\mathscr{B}((0, \infty]) \otimes \mathscr{P}(\{\mathrm{F}, \mathrm{G}\})), \tag{S5}$$

where $\mathscr{B}(S_1)$ denotes the Borel-$\sigma$-algebra generated from a uncountable set $S_1$, and $\mathscr{P}(S_2)$ denotes the power set generated from a countable set $S_2$. And, the corresponding probability measure $\mathbb{P}$ on $(\Omega, \mathscr{F})$ can be constructed recursively from the conditional transition probabilities by Ionescu-Tulcea's theorem [20].

For each path $\omega \in \Omega$, define the jump times $(\widetilde{t}_n)_{n \in \mathbb{N}_{\geq 0}}$ by

$$\widetilde{t}_n(\omega) := \begin{cases} \widetilde{t}_{n-1}(\omega) + s_{n-1}, & n > 0, \\ 0, & n = 0. \end{cases} \tag{S6}$$

And for every observation time $t \in \mathscr{T} = [0, \infty)$, let

$$N_t(\omega) = \max\{n \geq 0 : \widetilde{t}_n(\omega) \leq t\} \tag{S7}$$

be the number of reactions occurred by $t$. Then, the state of the system $(X_t, Y_t, Z_t)$ at $t$, described by the number of each particle type $(n_{S_0}(t), n_R(t), n_{S_R}(t))$, is obtained by starting from the initial state $(X_0, Y_0, Z_0) = (\alpha, \beta, 0)$ with $\alpha, \beta \in \mathbb{N}$ and applying sequentially the reactions $\xi_0, \ldots, \xi_{N_t(\omega)-1}$, where each reaction modifies the state according to

$$\begin{cases} (x, y, z) \mapsto (x-1, y-1, z+1), & \xi_n = \mathrm{F}, \\ (x, y, z) \mapsto (x+2, y, z-1), & \xi_n = \mathrm{G}. \end{cases} \tag{S8}$$

Thus, the state of the system for some fixed $t \in \mathscr{T}$ is determined entirely by the map

$$\omega \mapsto (X_t(\omega), Y_t(\omega), Z_t(\omega)), \tag{S9}$$

where the triple $(X_t, Y_t, Z_t) : \Omega \to \mathscr{S}$ is an $\mathscr{S}$-valued random variable that takes value from the finite, discrete state space

$$\mathscr{S} = \{(x, y, z) \in \mathbb{N}^3_{\geq 0} : x + y + 2z = \alpha + \beta\}. \tag{S10}$$

And $(X_t, Y_t, Z_t)_{t \in \mathscr{T}}$ describes a continuous-time Markov chain on $(\Omega, \mathscr{F}, \mathbb{P})$, with transition rates from the state $(x, y, z)$ being $kxy$ for feeding and $gz$ for growth. Physically, the trajectory traced by $(X_t, Y_t, Z_t)_{t \in \mathscr{T}}$ is a three-dimensional right-continuous function composed of a finite linear combination of scaled indicator functions, also known as a simple function.

Now, let the joint probability of a state be

$$P_{x,y,z}(t) = \begin{cases} \mathbb{P}[X_t = x, Y_t = y, Z_t = z], & (x, y, z) \in \mathscr{S}, \\ 0, & \text{otherwise.} \end{cases} \tag{S11}$$

As $z$ can be expressed in terms of $x, y$, we write $P_{x,y}(t) = P_{x,y,z}(t)$. Then we have

$$P_{x,y}(t+\Delta t) = k(x+1)(y+1)\Delta t P_{x+1,y+1}(t) + g(z+1)\Delta t P_{x-2,y}(t) + [1-(kxy+gz)\Delta t]P_{x,y}(t) + \mathscr{O}(\Delta t^2). \tag{S12}$$

Taking $\Delta t \searrow 0$ yields the chemical master equation

$$\frac{\mathrm{d}}{\mathrm{d}t} P_{x,y}(t) = k(x+1)(y+1)P_{x+1,y+1}(t) + g(z+1)P_{x-2,y}(t) - (kxy + gz)P_{x,y}(t). \tag{S13}$$

Monte Carlo simulation is used to study the stochastic kinetics of our coarse-grained system numerically exactly [21–24], which obeys the mechanism in (S1) consisting of two reaction channels. At each jump, the waiting time $\Delta t$ is drawn from an exponential distribution with the parameter $\lambda_0(t) = kX_tY_t + gZ_t$. The feeding reaction channel is selected with probability $kX_tY_t/\lambda_0(t)$ and the growth reaction channel with probability $gZ_t/\lambda_0(t)$, implemented by drawing a number uniformly on the interval $[0, \lambda_0(t)]$. The initial state is $(X_0, Y_0, Z_0) = (\alpha, \beta, 0)$, and the simulation runs until a specified maximum time or until $\lambda_0(t) = 0$.

All code files used to perform the analytical calculation, numerical simulation, and figure generation in this study are publicly available at `https://github.com/cfaleung1-rice/publication_1r1s` under the MIT license. The same repository can be found at `https://doi.org/10.5281/zenodo.21252284` [25].

### S2 Proof of Timescale Separation at Extreme Values of $k$ and $g$

The following statements enable us to systematically construct the scheme of limiting cases in Figure 2.

**Theorem S2.1.** *The following are true, $\mathbb{P}$-almost surely:*

1. *Feeding occurs given $X_tY_t > 0$ and $Z_t = 0$.*
2. *Growth occurs given $X_tY_t = 0$ and $Z_t > 0$.*

*Proof.* By the conservation law (S2), set $x + y + 2z = \alpha + \beta$. Then

$$\mathbb{P}[\xi_n = \mathrm{F} \mid X_{t_n} = x, Y_{t_n} = y, Z_{t_n} = z] = \frac{kxy}{kxy + gz}, \tag{S14}$$

$$\mathbb{P}[\xi_n = \mathrm{G} \mid X_{t_n} = x, Y_{t_n} = y, Z_{t_n} = z] = \frac{gz}{kxy + gz}. \tag{S15}$$

The results are trivial by direct substitution. □

**Corollary S2.1.1.** *If $g/k \nearrow \infty$, then growth occurs once $Z_t = 1$, with probability approaching 1.*

*Proof.* Denote the event as $\mathscr{E}_1$. By Theorem S2.1, we have

$$\begin{aligned}\mathbb{P}[\mathscr{E}_1] &= \prod_{n=1}^{\beta-1} \frac{g}{k(\alpha+n-2)(\beta-n)+g} \\ &= \prod_{n=1}^{\beta-1} \frac{1}{(g/k)^{-1}(\alpha+n-2)(\beta-n)+1} \\ &\xrightarrow{g/k\nearrow\infty} 1.\end{aligned} \tag{S16}$$

□

**Corollary S2.1.2.** *Define $\tau_{S_0}$ and $\tau_R$ by*

$$\tau_{S_0} := \inf\{t \in \mathscr{T} : X_t = 0\}, \tag{S17}$$

$$\tau_R := \inf\{t \in \mathscr{T} : Y_t = 0\}, \tag{S18}$$

*which we call the first extinction time of $X_t$ and $Y_t$, respectively. If $g/k \searrow 0$, then only feeding occurs for $t < \min\{\tau_{S_0}, \tau_R\}$, with probability approaching 1.*

*Proof.* Denote the event by $\mathscr{E}_2$. By Theorem S2.1, we have

$$\begin{aligned}\mathbb{P}[\mathscr{E}_2] &= \prod_{n=1}^{\min\{\alpha,\beta\}-1} \frac{k(\alpha-n)(\beta-n)}{k(\alpha-n)(\beta-n)+gn} \\ &= \prod_{n=1}^{\min\{\alpha,\beta\}-1} \frac{(\alpha-n)(\beta-n)}{(\alpha-n)(\beta-n)+(g/k)n} \\ &\xrightarrow{g/k\searrow 0} 1.\end{aligned} \tag{S19}$$

□

**Corollary S2.1.3.** *Suppose $g/k \searrow 0$. The following are true, with probability approaching 1:*

1. *If $\alpha < \beta$, then $\tau_{S_0} < \tau_R$.*
2. *If $\alpha = \beta$, then $\tau_{S_0} = \tau_R$.*

3. *If $\alpha > \beta$, then $\tau_{S_0} > \tau_R$, and $\tau_{S_0} = \infty$.*

*Proof.* The results are trivial and directly follow from Theorem S2.1 and Corollary S2.1.2. $\square$

**Corollary S2.1.4.** *Suppose $\alpha < \beta$. If $g/k \searrow 0$, then only feeding occurs for $\tau_{S_0} < t \leq \tau_R$ whenever $X_t > 0$, with probability approaching* 1.

*Proof.* Denote the event by $\mathscr{E}_3$. By definition, $X_{\tau_{S_0}} = 0$, so $Y_{\tau_{S_0}} = \beta - \alpha$ and $Z_{\tau_{S_0}} = \alpha$. Thus, growth must occur first. Suppose growing occurs at $\tau_{S_0} + \varepsilon$ where $\varepsilon > 0$. Then $X_{\tau_{S_0}+\varepsilon} = 2$, $Y_{\tau_{S_0}+\varepsilon} = \beta - \alpha$, and $Z_{\tau_{S_0}+\varepsilon} = \alpha - 1$. So, for a path on $\mathscr{E}_3$, two consecutive feeding reactions must occur. This implies the associated probability depends on the parity of $\beta - \alpha$.

Now, we can write

$$\mathbb{P}[\mathscr{E}_3] = \begin{cases} \displaystyle\prod_{n=0}^{(\beta-\alpha-2)/2} \prod_{m=1}^{2} Q(n,m), & 2 \mid (\beta - \alpha), \\ \displaystyle Q\left(\frac{\beta-\alpha-1}{2}, 2\right) \prod_{n=0}^{(\beta-\alpha-3)/2} \prod_{m=1}^{2} Q(n,m), & \text{otherwise.} \end{cases}$$
$$\xrightarrow{g/k \searrow 0} 1. \tag{S20}$$

where

$$\begin{aligned} Q(n,m) &:= \frac{km[\beta - \alpha - 2n + (m-2)]}{km[\beta - \alpha - 2n + (m-2)] + g[\alpha + n - (m-1)]} \\ &= \frac{m[\beta - \alpha - 2n + (m-2)]}{m[\beta - \alpha - 2n + (m-2)] + (g/k)[\alpha + n - (m-1)]}. \end{aligned} \tag{S21}$$

$\square$

These statements allow us to lump certain transitions together into simplified effective transitions when $g/k \nearrow \infty$ to $g/k \searrow 0$. From Fig. S2.1, it is easy to see how certain limiting cases are segregated into distinct temporal phases. For convenience, in later sections, we will denote the fast-growing limit as "A", the fast-feeding limit with $\alpha \geq \beta$ as "B", and the fast-feeding limit with $\alpha < \beta$ as "C". Temporal phases within each limiting regime are labelled by roman numerals from (i) up to (iii).

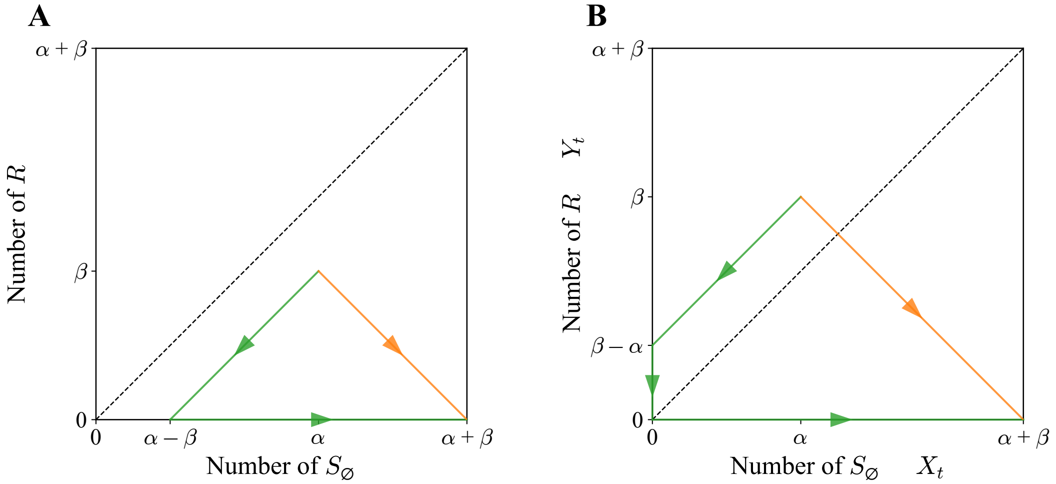

Figure S2.1: Trajectories of the single-species-single-resource system in the state space under different limiting conditions. (**A**) $\alpha \geq \beta$. (**B**) $\alpha < \beta$. Orange and green curves represent the $g/k \nearrow \infty$ and $g/k \searrow 0$ cases, respectively.

### S3 Solutions to the Coarse-Grained Forward Master Equation in the Fast-Growing Limit ($g/k \nearrow \infty$)

In this limit, there is no temporal phase transition. As $S_R$ is only transiently produced, we consider $Z_t = 0$ such that $X_t + Y_t = \alpha + \beta$. It suffices to only focus on $Y_t$ provided that the joint probability distribution $P^{(A)}_{x,y,z}(t)$ can be fully recovered from the single-variable case by

$$P^{(A)}_y(t) := P^{(A)}_{\alpha+\beta-y,y,0}(t), \tag{S22}$$

which is detailed in the Main Text. The evolution of $Y_t$ describes a pure-death process with discrete states $y \in \{0, \ldots, \beta\}$. Denote $y = 0$ as the absorbing state.

When $y = \beta$,

$$\begin{aligned} P^{(A)}_y(t+\Delta t) &= (1 - kxy\Delta t)P_y(t) \\ \Rightarrow \quad \frac{\mathrm{d}}{\mathrm{d}t}P_y(t) &= -k\alpha\beta P_y(t). \end{aligned} \tag{S23}$$

When $y \in \{0, ..., \beta - 1\}$,

$$\begin{aligned} P^{(A)}_y(t+\Delta t) &= k(x-1)(y+1)\Delta t P^{(A)}_{y+1}(t) + (1 - kxy\Delta t)P^{(A)}_y(t) + \mathscr{O}(\Delta t^2) \\ \Rightarrow \quad \frac{\mathrm{d}}{\mathrm{d}t}P^{(A)}_y(t) &= k(\alpha+\beta-y-1)(y+1)P^{(A)}_{y+1}(t) - k(\alpha+\beta-y)(y)P^{(A)}_y(t) \\ &= k[\alpha+\beta-(y+1)](y+1)P^{(A)}_{y+1}(t) - k(\alpha+\beta-y)(y)P^{(A)}_y(t). \end{aligned} \tag{S24}$$

Let $\lambda^{(A)}_i = k(\alpha+\beta-i)i$ for $i \geq 0$, then overall we have

$$\frac{\mathrm{d}}{\mathrm{d}t}P^{(A)}_y(t) = \begin{cases} -\lambda^{(A)}_y P_y(t), & y = \beta, \\ \lambda^{(A)}_{y+1}P_{y+1}(t) - \lambda^{(A)}_y P_y(t), & y \in \{0, ..., \beta-1\}. \end{cases} \tag{S25}$$

Using the initial condition

$$P^{(A)}_y(0) = \delta_{y,\beta} \tag{S26}$$

where $\delta$ denotes the Kronecker delta function, Laplace transform $\mathscr{L}\{\cdot\}(s)$ of the forward master equation yields

$$s\widetilde{P}^{(A)}_\beta(s) - \overbrace{P^{(A)}_\beta(0)}^{=1} = -\lambda^{(A)}_\beta \widetilde{P}^{(A)}_\beta(s) \quad \Rightarrow \quad \widetilde{P}^{(A)}_\beta(s) = \frac{1}{s+\lambda^{(A)}_\beta}, \tag{S27}$$

$$s\widetilde{P}^{(A)}_y(s) - \overbrace{P^{(A)}_y(0)}^{=0} = \lambda^{(A)}_{y+1}\widetilde{P}^{(A)}_{y+1}(s) - \lambda^{(A)}_y \widetilde{P}^{(A)}_y(s) \quad \Rightarrow \quad \widetilde{P}^{(A)}_y(s) = \frac{\lambda^{(A)}_{y+1}}{s+\lambda^{(A)}_y}\widetilde{P}^{(A)}_{y+1}(s), \quad y < \beta, \tag{S28}$$

where $\widetilde{P}$ denotes the Laplace-transformed distribution.

Thus, we have

$$\widetilde{P}^{(A)}_y(s) = \frac{\lambda^{(A)}_{y+1}}{s+\lambda^{(A)}_y} \cdots \frac{\lambda^{(A)}_\beta}{s+\lambda^{(A)}_{\beta-1}} \frac{1}{s+\lambda^{(A)}_\beta} = \frac{1}{\lambda^{(A)}_y}\prod_{n=y}^{\beta} \frac{\lambda^{(A)}_n}{s+\lambda^{(A)}_n}, \quad y > 0. \tag{S29}$$

To perform the inverse Laplace transform $\mathscr{L}^{-1}\{\cdot\}(t)$, we decompose the product into partial fractions. Set $y > 0$. Because $\lambda^{(A)}_y = ky(\alpha+\beta-y)$, the symmetry $\lambda^{(A)}_y = \lambda^{(A)}_{\alpha+\beta-y}$ means that degenerate poles may exist depending on the $y$ chosen. Let $\mathscr{I}_y = \{n \in \mathbb{N} : y \leq n \leq \beta\}$ be the set of indices appearing in the product. Denote the distinct values of $\lambda^{(A)}_n$ for $n \in \mathscr{I}_y$ by $\zeta_1, \ldots, \zeta_{K_y}$ with $\zeta_a < \zeta_b$ for all $a < b$, where

$K_y = |\{\lambda_n^{(\mathrm{A})} : n \in \mathscr{I}_y\}|$ and $|\cdot|$ denotes the cardinality of a set. For each distinct pole $\zeta_j$, its multiplicity in the product is

$$W_j = |\{n \in \mathscr{I}_y : \lambda_n = \zeta_j\}| \,. \tag{S30}$$

In particular, if $\alpha+\beta$ is even and $(\alpha+\beta)/2 \in \mathscr{I}_y$, then its corresponding pole has $W_j = 1$. And if there exists some $n \in \mathscr{I}_y$ such that $\alpha + \beta - n \in \mathscr{I}_y$, then its corresponding pole has $W_j = 2$. This means $W_j \in \{1, 2\}$. So, the partial fraction expansion reads

$$\prod_{n=y}^{\beta} \frac{\lambda_n^{(\mathrm{A})}}{s + \lambda_n^{(\mathrm{A})}} = \sum_{n=1}^{K_y} \sum_{m=1}^{W_n} \frac{A_{n,m}}{(s + \zeta_n)^m}, \tag{S31}$$

where $A_{n,m}$ has the well-known form [26]:

$$A_{n,m} = \frac{1}{(W_n - m)!} \left. \frac{\mathrm{d}^{W_n - m}}{\mathrm{d}s^{W_n - m}} \phi_n(s) \right|_{s=-\zeta_n}, \tag{S32}$$

$$\phi_n(s) = \frac{\prod_{i=y}^{\beta} \lambda_i^{(\mathrm{A})}}{\prod_{\substack{j=y, \\ \lambda_j^{(\mathrm{A})} \neq \zeta_n}}^{\beta} (s + \lambda_j^{(\mathrm{A})})}. \tag{S33}$$

Thus, we have

$$\begin{aligned} P_y^{(\mathrm{A})}(t) &= \frac{1}{\lambda_y^{(\mathrm{A})}} \sum_{n=1}^{K_y} \sum_{m=1}^{W_n} A_{n,m} \mathscr{L}^{-1} \left\{ (s + \zeta_n)^{-m} \right\} (t) \\ &= \frac{1}{\lambda_y^{(\mathrm{A})}} \sum_{n=1}^{K_y} \sum_{m=1}^{W_n} \frac{A_{n,m}}{(m-1)!} t^{m-1} e^{-\zeta_n t}, \quad y > 0. \end{aligned} \tag{S34}$$

For $y = 0$, (S28) allows us to write

$$\begin{aligned} P_0^{(\mathrm{A})}(t) &= \mathscr{L}^{-1} \left\{ \frac{\lambda_1^{(\mathrm{A})}}{s} \widetilde{P}_1^{(\mathrm{A})}(s) \right\} (t) \\ &= \lambda_1^{(\mathrm{A})} \int_0^t P_1^{(\mathrm{A})}(u) \, \mathrm{d}u \\ &= \lambda_1^{(\mathrm{A})} \overbrace{\left[ \frac{1}{\lambda_1^{(\mathrm{A})}} \sum_{n=1}^{K_1} \sum_{m=1}^{W_n} \frac{A_{n,m}}{(m-1)!} \int_0^t u^{m-1} e^{-\zeta_n u} \, \mathrm{d}u \right]}^{\text{exchange } \int \text{ and } \sum \text{ by Fubini-Tonelli}} \\ &= \sum_{n=1}^{K_1} \sum_{m=1}^{W_n} \frac{A_{n,m}}{(m-1)!} \frac{1}{\zeta_n^m} \overbrace{\int_0^{\zeta_n t} (\zeta_n u)^{m-1} e^{-\zeta_n u} \, \mathrm{d}(\zeta_n u)}^{=:\gamma(m, \zeta_n t)} \\ &= \sum_{n=1}^{K_1} \sum_{m=1}^{W_n} \frac{A_{n,m}}{(m-1)!} \frac{1}{\zeta_n^m} \left[ (\zeta_n t)^m e^{-\zeta_n t} \sum_{\ell=0}^{\infty} \frac{(m-1)!}{(m+\ell)!} (\zeta_n t)^\ell \right] \\ &= \sum_{n=1}^{K_1} \sum_{m=1}^{W_n} A_{n,m} t^m e^{-\zeta_n t} \sum_{\ell=0}^{\infty} \frac{(\zeta_n t)^\ell}{(m+\ell)!}, \end{aligned} \tag{S35}$$

where $\gamma$ is the lower incomplete gamma function [27]. The moments $\mathbb{E}^{(\mathrm{A})}[X_t^n], \mathbb{E}^{(\mathrm{A})}[Y_t^n], \mathbb{E}^{(\mathrm{A})}[Z_t^n]$ can then be computed by definition, for $n \in \mathbb{N}$. Recall that we take the convention which $P_y^{(\mathrm{A})}(t) = 0$ if $y \notin \{0, \ldots, \beta\}$ [28], such that we can evaluate the expectations as infinite sums in the Main Text.

### S4 Solutions to the Coarse-Grained Forward Master Equation in the Resource-Limited Fast-Feeding Limit ($g/k \searrow 0, \alpha \geq \beta$), in the First Temporal Phase

In this limit, there is one temporal phase transition. In the first phase, $R$ is rapidly depleted. As $S_0$ combines with $R$ in equal parts, $X_t = \alpha - \beta + Y_t$ such that $(\alpha - \beta + Y_t) + Y_t + 2Z_t = \alpha + \beta \Rightarrow Z_t = \beta - Y_t$, and

$$P_y^{(\mathrm{B,i})}(t) := P_{\alpha-\beta+y,y,\beta-y}(t). \tag{S36}$$

The evolution of $Y_t$ is this phase describes a pure-death process with discrete states $y \in \{0,\ldots,\beta\}$. Denote $y = 0$ as the absorbing state.

When $y = \beta$,

$$\begin{aligned} P_y^{(\mathrm{B,i})}(t+\Delta t) &= (1 - kxy\Delta t)P_y^{(\mathrm{B,i})}(t) \\ \Rightarrow \quad \frac{\mathrm{d}}{\mathrm{d}t}P_y^{(\mathrm{B,i})}(t) &= -k\alpha\beta P_y^{(\mathrm{B,i})}(t). \end{aligned} \tag{S37}$$

When $y \in \{1,...,\beta-1\}$,

$$\begin{aligned} P_y^{(\mathrm{B,i})}(t+\Delta t) &= k(x+1)(y+1)\Delta t P_{y+1}^{(\mathrm{B,i})}(t) + (1 - kxy\Delta t)P_y^{(\mathrm{B,i})}(t) + \mathcal{O}(\Delta t^2) \\ \Rightarrow \quad \frac{\mathrm{d}}{\mathrm{d}t}P_y^{(\mathrm{B,i})}(t) &= k[\alpha - \beta + (y+1)](y+1)P_{y+1}^{(\mathrm{B,i})}(t) - k(\alpha-\beta+y)yP_y^{(\mathrm{B,i})}(t). \end{aligned} \tag{S38}$$

When $y = 0$,

$$\begin{aligned} P_y^{(\mathrm{B,i})}(t+\Delta t) &= k(x+1)(y+1)\Delta t P_{y+1}^{(\mathrm{B,i})}(t) + P_y^{(\mathrm{B,i})}(t) + \mathcal{O}(\Delta t^2) \\ \Rightarrow \quad \frac{\mathrm{d}}{\mathrm{d}t}P_y^{(\mathrm{B,i})}(t) &= k(\alpha-\beta+1)P_{y+1}^{(\mathrm{B,i})}(t). \end{aligned} \tag{S39}$$

Let $\lambda_i^{(\mathrm{B,i})} = k(\alpha - \beta + i)i$ for $i \geq 0$, then overall we have

$$\frac{\mathrm{d}}{\mathrm{d}t}P_y^{(\mathrm{B,i})}(t) = \begin{cases} -\lambda_y^{(\mathrm{B,i})}P_y^{(\mathrm{B,i})}(t), & y = \beta, \\ \lambda_{y+1}^{(\mathrm{B,i})}P_{y+1}^{(\mathrm{B,i})}(t) - \lambda_y^{(\mathrm{B,i})}P_y^{(\mathrm{B,i})}(t), & y \in \{0,...,\beta-1\}. \end{cases} \tag{S40}$$

The functional form of (S40) is virtually identical to (S25), so their solutions are almost the same only with different transition probabilities. However, notice $\lambda_y^{(\mathrm{B,i})},\ldots,\lambda_\beta^{(\mathrm{B,i})}$ are mutually distinct for any $y$, meaning the partial fraction expression can be simplified further.

Let $\mathcal{Y}^{(\mathrm{B,i})} = \{0,\ldots,\beta\}$ and define $\mathcal{Y}_y^{(\mathrm{I})} := \{n \in \mathcal{Y}^{(\mathrm{I})} : n \geq y\}$ for some temporal phase I. Then, in the $s$-domain, we can do the partial fraction decomposition

$$\begin{aligned} \prod_{n=y}^{\beta} \frac{\lambda_n^{(\mathrm{B,i})}}{s + \lambda_n^{(\mathrm{B,i})}} &= \sum_{n=y}^{\beta} \frac{\kappa_n^{(\mathrm{B,i})}}{s + \lambda_n} \\ \Rightarrow \quad \prod_{n=y}^{\beta} \lambda_n^{(\mathrm{B,i})} &= \sum_{n=y}^{\beta} \kappa_n^{(\mathrm{B,i})} \prod_{m \in \mathcal{Y}_y^{(\mathrm{B,i})} \setminus \{n\}} (s + \lambda_m^{(\mathrm{B,i})}). \end{aligned} \tag{S41}$$

Whilst the former fractional equation holds only for $s$ such that $s + \lambda_n^{(\mathrm{B,i})} \neq 0$ for $n \in \{y,\ldots,\beta\}$, the latter polynomial identity holds for all $s$. Choose $s = -\lambda_j^{(\mathrm{B,i})}$ for some fixed $j \in \{y,\ldots,\beta\}$, then all terms of the sum except for $n = j$ vanish:

$$\prod_{n=y}^{\beta} \lambda_n^{(\mathrm{B,i})} = \kappa_j^{(\mathrm{B,i})} \prod_{m \in \mathscr{Y}_y^{(\mathrm{B,i})} \setminus \{j\}} (\lambda_m^{(\mathrm{B,i})} - \lambda_j^{(\mathrm{B,i})})$$

$$\Rightarrow \quad \kappa_j^{(\mathrm{B,i})} = \frac{\prod_{n=y}^{\beta} \lambda_n^{(\mathrm{B,i})}}{\prod_{m \in \mathscr{Y}_y^{(\mathrm{B,i})} \setminus \{j\}} (\lambda_m^{(\mathrm{B,i})} - \lambda_j^{(\mathrm{B,i})})}. \tag{S42}$$

Note that $\kappa_j^{(\mathrm{B,i})}$ implicitly depends on $y$, and we take the empty product convention if $\mathscr{Y}_y^{(\mathrm{B,i})} \setminus \{j\} = \varnothing$. Thus, we have

$$\begin{aligned} P_y^{(\mathrm{B,i})}(t) &= \frac{1}{\lambda_y^{(\mathrm{B,i})}} \sum_{n=y}^{\beta} \kappa_n^{(\mathrm{B,i})} \mathscr{L}^{-1} \left\{ \frac{1}{s - (-\lambda_n^{(\mathrm{B,i})})} \right\} (t) \\ &= \frac{1}{\lambda_y^{(\mathrm{B,i})}} \sum_{n=y}^{\beta} \kappa_n^{(\mathrm{B,i})} e^{-\lambda_n^{(\mathrm{B,i})} t}, \quad y > 0. \end{aligned} \tag{S43}$$

For $y = 0$, we follow (S35) to get

$$\begin{aligned} P_0^{(\mathrm{B,i})}(t) &= \lambda_1^{(\mathrm{B,i})} \int_0^t P_1^{(\mathrm{B,i})}(u)\,\mathrm{d}u \\ &= \sum_{n=1}^{\beta} \kappa_n^{(\mathrm{B,i})} \int_0^t e^{-\lambda_n^{(\mathrm{B,i})} u}\,\mathrm{d}u \\ &= \sum_{n=1}^{\beta} \frac{\kappa_n^{(\mathrm{B,i})}}{\lambda_n^{(\mathrm{B,i})}} (1 - e^{-\lambda_n^{(\mathrm{B,i})} t}). \end{aligned} \tag{S44}$$

Solutions to the coarse-grained forward master equations associated to the second temporal phase is well-known [28, 29], and can be found in the Main Text.

### S5 Moments of the Random Variables in the Resource-Limited Fast-Feeding Limit ($g/k \searrow 0, \alpha \geq \beta$)

Take

$$P_y^{(\mathrm{B,i})}(t) = \begin{cases} \dfrac{1}{\lambda_y^{(\mathrm{B,i})}} \displaystyle\sum_{n=y}^{\beta} \kappa_n^{(\mathrm{B,i})} e^{-\lambda_n^{(\mathrm{B,i})} t}, & y \in \{1, \dots, \beta\}, \\ \displaystyle\sum_{n=1}^{\beta} \frac{\kappa_n^{(\mathrm{B,i})}}{\lambda_n^{(\mathrm{B,i})}} (1 - e^{-\lambda_n^{(\mathrm{B,i})} t}), & y = 0. \end{cases} \tag{S45}$$

Also, take

$$P_z^{(\mathrm{B,ii})}(t) = \binom{\beta}{z} (e^{-gt})^z (1 - e^{-gt})^{\beta - z}. \tag{S46}$$

With $\mathscr{T} = [0, \infty)$, consider the Markov chains $(X_t)_{t\in\mathscr{T}}, (Y_t)_{t\in\mathscr{T}}, (Z_t)_{t\in\mathscr{T}}$ on $(\Omega, \mathscr{F})$ with the laws $\mathbb{P}_{X_t}, \mathbb{P}_{Y_t}, \mathbb{P}_{Z_t}$ respectively, which have densities $f_{X_t}, f_{Y_t}, f_{Z_t}$ absolutely continuous with the counting measure $\mu_{\mathbb{Z}}$. These are projections of the stochastic process $(X_t, Y_t, Z_t)_{t\in\mathscr{T}}$ introduced in the Main Text. Note that the mean first extinction times $\tau_{S_0}, \tau_R$ are also random variables on $(\Omega, \mathscr{F})$. Denote the densities of $\tau_{S_0}, \tau_R$ by $f_{\tau_{S_0}}, f_{\tau_R}$ respectively, which are absolutely continuous with the one-dimensional Lebesgue measure.

Then, by the law of iterated expectations, for some fixed observation time $t \in \mathscr{T}$, we can write the $n^{\text{th}}$ moment of $Z_t$ as

$$\begin{aligned}
\mathbb{E}^{(\mathrm{B})}[Z_t^n] &= \mathbb{E}_{\tau_R}[\mathbb{E}[Z_t^n | \tau_R]] \\
&= \int_0^\infty \mathbb{E}[Z_t^n | \tau_R = u] f_{\tau_R}(u)\,\mathrm{d}u \\
&= \overbrace{\int_0^t \mathbb{E}[Z_t^n | \tau_R = u] f_{\tau_R}(u)\,\mathrm{d}u}^{\substack{\text{post-extinction of } R: \\ \tau_R = u \leq t,\ R = 0 \text{ at } t}} + \overbrace{\int_t^\infty \mathbb{E}[Z_t^n | \tau_R = u] f_{\tau_R}(u)\,\mathrm{d}u}^{\substack{\text{pre-extinction of } R: \\ \tau_R = u > t,\ R > 0 \text{ at } t}} \\
&= \int_0^\infty \mathbb{E}[Z_t^n | \tau_R = u] \mathbb{1}_{[0,t]} f_{\tau_R}(u)\,\mathrm{d}u + \int_0^\infty \mathbb{E}[Z_t^n | \tau_R = u] \mathbb{1}_{(t,\infty)} f_{\tau_R}(u)\,\mathrm{d}u \\
&= \mathbb{E}_{\tau_R}[\mathbb{E}[Z_t^n|\tau_R] \mathbb{1}_{\{\tau_R \leq t\}}] + \mathbb{E}_{\tau_R}[\mathbb{E}[Z_t^n|\tau_R]\, \mathbb{1}_{\{\tau_R > t\}}] \\
&= \mathbb{E}[Z_t^n \mathbb{1}_{\{\tau_R \leq t\}}] + \mathbb{E}[Z_t^n \mathbb{1}_{\{\tau_R > t\}}] \\
&= \overbrace{\mathbb{E}[Z_t^n \mathbb{1}_{\{\tau_R > t\}}]}^{\text{pre-}\tau_R} + \overbrace{\mathbb{E}[Z_t^n \mathbb{1}_{\{\tau_R \leq t\}}]}^{\text{post-}\tau_R}.
\end{aligned} \tag{S47}$$

In the pre-$\tau_R$ phase, $\{\tau_R > t\} = \{Y_t > 0\}$ almost surely. So, $\mathbb{E}[Z_t^n \mathbb{1}_{\{\tau_R > t\}}] = \mathbb{E}[(\beta - Y_t)^n \mathbb{1}_{\{\tau_R > t\}}]$. Hence,

$$\begin{aligned}
\mathbb{E}[Z_t^n \mathbb{1}_{\{\tau_R > t\}}] &= \mathbb{E}\left[(\beta - Y_t)^n \mathbb{1}_{\{Y_t > 0\}}\right] \\
&= \int_\Omega (\beta - Y_t)^n \mathbb{1}_{\{Y_t > 0\}}\,\mathrm{d}\mathbb{P}_{Y_t}(\omega) \\
&= \int_{(0,\infty]} (\beta - y)^n f_{Y_t}^{(\mathrm{B,i})}(y)\,\mathrm{d}\mu_{\mathbb{Z}}(y) \\
&= \sum_{y=1}^{\infty} (\beta - y)^n P_y^{(\mathrm{B,i})}(t)
\end{aligned} \tag{S48}$$

In the post-$\tau_R$ phase, $\{\tau_R \leq t\} = \{Y_t = 0\}$ almost surely, so we cannot use the same approach as (S48).

Instead, notice $\mathbb{E}[Z_t^n|\tau_R = u] = \mathbb{E}^{(\mathrm{B},i)}[Z_{t-u}^n]$ for any $\tau_R = u \le t$ by the strong Markov property. Hence,

$$\begin{aligned}
\mathbb{E}[Z_t^n \mathbb{1}_{\{\tau_R \le t\}}] &= \int_\Omega \mathbb{E}[Z_t^n|\tau_R]\mathbb{1}_{\{\tau_R \le t\}}\,\mathrm{d}\mathbb{P}_{\tau_R}(\omega) \\
&= \int_{[0,t]} \mathbb{E}^{(\mathrm{B},i)}[Z_{t-u}^n] f_{\tau_R}(u)\,\mathrm{d}u \\
&= \int_0^t \sum_{z=0}^{\infty} z^n P_z^{(\mathrm{B,ii})}(t-u) f_{\tau_R}(u)\,\mathrm{d}u, \qquad \text{(S49)}
\end{aligned}$$

where

$$\begin{aligned}
f_{\tau_R}(u) &:= \frac{\mathrm{d}}{\mathrm{d}u}\mathbb{P}[\tau_R \le u] \\
&= \frac{\mathrm{d}}{\mathrm{d}u}\mathbb{P}[Y_u = 0] \\
&= \sum_{n=1}^{\beta} \kappa_n^{(\mathrm{B,i})} e^{-\lambda_n^{(\mathrm{B,i})} u}. \qquad \text{(S50)}
\end{aligned}$$

Accordingly, we have

$$\mathbb{E}^{(\mathrm{B})}[X_t^n] = \sum_{y=1}^{\infty} (\alpha - \beta + y)^n P_y^{(\mathrm{B,i})}(t) + \int_0^t \left[\sum_{z=0}^{\infty} (\alpha + \beta - 2z)^n P_z^{(\mathrm{B,ii})}(t-u)\right] f_{\tau_R}(u)\,\mathrm{d}u, \qquad \text{(S51)}$$

$$\mathbb{E}^{(\mathrm{B})}[Y_t^n] = \sum_{y=1}^{\infty} y^n P_y^{(\mathrm{B,i})}(t), \qquad \text{(S52)}$$

$$\mathbb{E}^{(\mathrm{B})}[Z_t^n] = \sum_{y=1}^{\infty} (\beta - y)^n P_y^{(\mathrm{B,i})}(t) + \int_0^t \left(\sum_{z=0}^{\infty} z^n P_z^{(\mathrm{B,ii})}(t-u)\right) f_{\tau_R}(u)\,\mathrm{d}u. \qquad \text{(S53)}$$

The convolution term can be evaluated explicitly as well. Consider for $k_1, k_2 \in \mathbb{C}$ that

$$\begin{aligned}
\int_0^t e^{-k_1(t-u)} e^{-k_2 u}\,\mathrm{d}u &= e^{-k_1 t}\int_0^t e^{(k_1-k_2)u}\,\mathrm{d}u \\
&= \begin{cases} t e^{-k_1 t}, & k_1 = k_2, \\ \dfrac{e^{-k_1 t}}{k_1 - k_2}\left[e^{(k_1-k_2)t} - 1\right], & \text{otherwise}, \end{cases} \\
&=: h_t(k_1, k_2). \qquad \text{(S54)}
\end{aligned}$$

Also, let $L : z \mapsto mz + b$ be a linear function of $z$ where $m, b \in \mathbb{R}$. Then, by binomial expansion, we have

$$\begin{aligned}
&\int_0^t \sum_{z=0}^{\beta} (L(z))^n P_z^{(\mathrm{B,ii})}(t-u) f_{\tau_R}(u)\,\mathrm{d}u \\
&= \int_0^t \sum_{z=0}^{\beta} (L(z))^n \left\{\binom{\beta}{z}[e^{-g(t-u)}]^z \sum_{q=0}^{\beta-z}\binom{\beta-z}{q}[-e^{-g(t-u)}]^q\right\} \sum_{r=1}^{\beta} \kappa_r^{(\mathrm{B,i})} e^{-\lambda_r^{(\mathrm{B,i})} u}\,\mathrm{d}u \\
&= \sum_{z=0}^{\beta} (L(z))^n \binom{\beta}{z} \sum_{q=0}^{\beta-z}\sum_{r=1}^{\beta} \binom{\beta-z}{q}(-1)^q \kappa_r^{(\mathrm{B,i})} \int_0^t e^{-g(z+q)(t-u)} e^{-\lambda_r^{(\mathrm{B,i})} u}\,\mathrm{d}u \\
&= \sum_{z=0}^{\beta} (L(z))^n \binom{\beta}{z} \sum_{q=0}^{\beta-z}\sum_{r=1}^{\beta} \binom{\beta-z}{q}(-1)^q \kappa_r^{(\mathrm{B,i})} h_t\left(g(z+q), \lambda_r^{(\mathrm{B,i})}\right). \qquad \text{(S55)}
\end{aligned}$$

### S6 Solutions to the Coarse-Grained Forward Master Equation in the Species-Limited Fast-Feeding Limit ($g/k \searrow 0, \alpha < \beta$)

In this limit, there are two temporal phase transitions. In the first phase, $S_0$ is rapidly depleted, and the associated solution $P_y^{(\mathrm{C,i})}$ is analogous to (S45), but with $\lambda_i^{(\mathrm{C,i})} = k(\alpha - \beta + i)i$ for $i \geq \beta - \alpha$:

$$P_y^{(\mathrm{C,i})}(t) = \begin{cases} \dfrac{1}{\lambda_y^{(\mathrm{C,i})}} \displaystyle\sum_{n=y}^{\beta} \kappa_n^{(\mathrm{C,i})} e^{-\lambda_n^{(\mathrm{C,i})} t}, & y \in \{\beta - \alpha + 1, \dots, \beta\}, \\ \displaystyle\sum_{n=\beta-\alpha+1}^{\beta} \dfrac{\kappa_n^{(\mathrm{C,i})}}{\lambda_n^{(\mathrm{C,i})}} (1 - e^{-\lambda_n^{(\mathrm{C,i})} t}), & y = \beta - \alpha. \end{cases} \tag{S56}$$

In the second phase, $S_R$ splits into a pair of $S_0$, which are rapidly depleted again until $R$ runs out. Following Corollary S2.1.4, we consider $X_t = 0$ such that $Y_t + 2Z_t = \alpha + \beta$. This means the states which $Y_t$ assumes depend on the parity of $\beta - \alpha$. Mathematically, we have

$$P_y^{(\mathrm{C,ii})}(t) := \begin{cases} P^{(\mathrm{C,ii})}_{0,y,(\alpha+\beta-y)/2}(t), & 2 \mid (\beta - \alpha) \text{ or } y \neq 0, \\ P^{(\mathrm{C,ii})}_{1,0,(\alpha+\beta-1)/2}(t), & \text{otherwise}, \end{cases} \tag{S57}$$

with

$$y \in \mathscr{Y}^{(\mathrm{C,ii})} := \begin{cases} \{0, 2, 4, \dots, \beta - \alpha\}, & 2 \mid (\beta - \alpha), \\ \{0, 1, 3, \dots, \beta - \alpha\}, & \text{otherwise}. \end{cases} \tag{S58}$$

More specifically, this means if $\beta - \alpha$ is odd, then the last step occurs asymmetrically, in the sense that only one of the newly generated $S_0$ would consume the single remaining $R$ particle and generate an $S_R$, after which phase transition proceeds. But otherwise, we can reasonably coarse-grain the rapid sequential consumption of $R$ into a single step.

When $y = \beta - \alpha$,

$$\begin{aligned} P_y^{(\mathrm{C,ii})}(t + \Delta t) &= (1 - gz\Delta t) P_y^{(\mathrm{C,ii})}(t) \\ \Rightarrow \quad \frac{\mathrm{d}}{\mathrm{d}t} P_y^{(\mathrm{C,ii})}(t) &= -g\alpha P_y^{(\mathrm{C,ii})}(t). \end{aligned} \tag{S59}$$

When $y \in \mathscr{Y}^{(\mathrm{C,ii})} \setminus \{0, \beta - \alpha\}$,

$$\begin{aligned} P_y^{(\mathrm{C,ii})}(t + \Delta t) &= g(z-1)\Delta t P_{y+2}^{(\mathrm{C,ii})}(t) + (1 - gz\Delta t) P_y^{(\mathrm{C,ii})}(t) + \mathscr{O}(\Delta t^2) \\ \Rightarrow \quad \frac{\mathrm{d}}{\mathrm{d}t} P_y^{(\mathrm{C,ii})}(t) &= g\left(\frac{\alpha + \beta - y}{2} - 1\right) P_{y+2}^{(\mathrm{C,ii})}(t) - g\left(\frac{\alpha + \beta - y}{2}\right) P_y^{(\mathrm{C,ii})}(t) \\ &= \frac{g}{2}[\alpha + \beta - (y+2)] P_{y+2}^{(\mathrm{C,ii})}(t) - \frac{g}{2}(\alpha + \beta - y) P_y^{(\mathrm{C,ii})}(t). \end{aligned} \tag{S60}$$

When $y = 0$,

$$\begin{aligned} P_y^{(\mathrm{C,ii})}(t + \Delta t) &= \begin{cases} g(z-1)\Delta t P_{y+2}^{(\mathrm{C,ii})}(t) + \mathscr{O}(\Delta t^2), & 2 \mid (\beta - \alpha), \\ gz'\Delta t P_{y+1}^{(\mathrm{C,ii})}(t) + \mathscr{O}(\Delta t^2), & \text{otherwise}, \end{cases} \\ \Rightarrow \quad \frac{\mathrm{d}}{\mathrm{d}t} P_y^{(\mathrm{C,ii})}(t) &= \begin{cases} \dfrac{g}{2}(\alpha + \beta - 2) P_{y+2}^{(\mathrm{C,ii})}(t), & 2 \mid (\beta - \alpha), \\ \dfrac{g}{2}(\alpha + \beta - 1) P_{y+1}^{(\mathrm{C,ii})}(t), & \text{otherwise}, \end{cases} \end{aligned} \tag{S61}$$

where $z' := z_{y+1}$ due to asymmetric feeding, by considering $z_y = (\alpha + \beta - y)/2$.

Let $\lambda_i^{(\mathrm{C,ii})} = g(\alpha+\beta-i)/2$ for $i \geq 1$. Also, define $\mathscr{Y}_{a,b}^{(\mathrm{I})} := \{n \in \mathscr{Y}^{(\mathrm{I})} : a < n < b\}$ for some temporal phase I. Then overall we have

$$\frac{\mathrm{d}}{\mathrm{d}t} P_y^{(\mathrm{C,ii})}(t) = \begin{cases} -\lambda_y^{(\mathrm{C,ii})} P_y^{(\mathrm{C,ii})}(t), & y = \beta-\alpha, \\ \lambda_{y+2}^{(\mathrm{C,ii})} P_{y+2}^{(\mathrm{C,ii})}(t) - \lambda_y^{(\mathrm{C,ii})} P_y^{(\mathrm{C,ii})}(t), & y \in \mathscr{Y}_{0,\beta-\alpha}^{(\mathrm{C,ii})}, \\ \lambda_{y+2}^{(\mathrm{C,ii})} P_{y+2}^{(\mathrm{C,ii})}(t), & 2 \mid (\beta-\alpha) \text{ and } y = 0, \\ \lambda_{y+1}^{(\mathrm{C,ii})} P_{y+1}^{(\mathrm{C,ii})}(t), & 2 \nmid (\beta-\alpha) \text{ and } y = 0. \end{cases} \tag{S62}$$

The functional form of (S62) is virtually identical to (S40), so their solutions are almost the same only with different transition probabilities and step sizes. Thus,

$$P_y^{(\mathrm{C,ii})}(t) = \begin{cases} \dfrac{1}{\lambda_y^{(\mathrm{C,ii})}} \displaystyle\sum_{n \in \mathscr{Y}_y^{(\mathrm{C,ii})}} \kappa_n^{(\mathrm{C,ii})} e^{-\lambda_n^{(\mathrm{C,ii})} t}, & y \in \mathscr{Y}_1^{(\mathrm{C,ii})}, \\ \displaystyle\sum_{n \in \mathscr{Y}_2^{(\mathrm{C,ii})}} \frac{\kappa_n^{(\mathrm{C,ii})}}{\lambda_n^{(\mathrm{C,ii})}} (1 - e^{-\lambda_n^{(\mathrm{C,ii})} t}), & 2 \mid (\beta-\alpha) \text{ and } y = 0, \\ \displaystyle\sum_{n \in \mathscr{Y}_1^{(\mathrm{C,ii})}} \frac{\kappa_n^{(\mathrm{C,ii})}}{\lambda_n^{(\mathrm{C,ii})}} (1 - e^{-\lambda_n^{(\mathrm{C,ii})} t}), & 2 \nmid (\beta-\alpha) \text{ and } y = 0, \end{cases} \tag{S63}$$

where

$$\kappa_j^{(\mathrm{C,ii})} = \frac{\displaystyle\prod_{n \in \mathscr{Y}_y^{(\mathrm{C,ii})}} \lambda_n^{(\mathrm{C,ii})}}{\displaystyle\prod_{m \in \mathscr{Y}_y^{(\mathrm{C,ii})} \setminus \{j\}} (\lambda_m^{(\mathrm{C,ii})} - \lambda_j^{(\mathrm{C,ii})})} \tag{S64}$$

for $j \in \mathscr{Y}_y^{(\mathrm{C,ii})}$ for some fixed $y \in \mathscr{Y}^{(\mathrm{C,ii})}$.

In the third phase, parity of $\beta - \alpha$ likewise results in two mutually exclusive state spaces for $Z_t$:

$$P_z^{(\mathrm{C,iii})}(t) := \begin{cases} P_{\alpha+\beta-2z,0,z}^{(\mathrm{C,iii})}(t), & 2 \mid (\beta-\alpha) \text{ or } z \neq \dfrac{\alpha+\beta-1}{2}, \\ P_{1,0,(\alpha+\beta-1)/2}^{(\mathrm{C,iii})}(t), & \text{otherwise}, \end{cases} \tag{S65}$$

with

$$z \in \begin{cases} \left\{0, 1, 2, \ldots, \dfrac{\alpha+\beta}{2}\right\}, & 2 \mid (\beta-\alpha), \\ \left\{0, 1, 2, \ldots, \dfrac{\alpha+\beta-1}{2}\right\}, & \text{otherwise}, \end{cases} \tag{S66}$$

such that

$$P_z^{(\mathrm{C,iii})}(t) = \binom{\lfloor(\alpha+\beta)/2\rfloor}{z} (e^{-gt})^z (1 - e^{-gt})^{\lfloor(\alpha+\beta)/2\rfloor - z}. \tag{S67}$$

### S7 Moments of the Random Variables in the Species-Limited Fast-Feeding Limit ($g/k \searrow 0, \alpha < \beta$)

Take (S56), (S63), and (S67). Recall Corollary S2.1.3 which states that $\tau_{S_0} < \tau_R$ if $g/k \searrow 0$ and $\alpha < \beta$. Define the positive random variable

$$D := \tau_R - \tau_{S_0} > 0. \tag{S68}$$

Then, by the law of iterated expectations, for some fixed observation time $t \in \mathscr{T}$, we can write the $n^{\text{th}}$ moment of $Z_t$ as

$$\begin{aligned}
\mathbb{E}[Z_t^m] &= \mathbb{E}_{\tau_{S_0},\tau_R}[\mathbb{E}[Z_t^m|\tau_{S_0},\tau_R]] \\
&= \int_0^\infty \int_0^\infty \mathbb{E}[Z_t^m|\tau_{S_0} = u, \tau_R = v] f_{\tau_{S_0},\tau_R}(u,v)\,\mathrm{d}u\,\mathrm{d}v \\
&= \int_0^\infty \int_0^\infty \mathbb{E}[Z_t^m|\tau_{S_0} = u, \tau_R = v] f_{\tau_{S_0}}(u) f_{\tau_R|\tau_{S_0}}(v|u)\,\mathrm{d}u\,\mathrm{d}v \\
&\xrightarrow{g/k\searrow 0} \int_0^\infty \int_0^\infty \mathbb{E}[Z_t^m|\tau_{S_0} = u, \tau_R = v] f_{\tau_{S_0}}(u) f_D(v-u)\,\mathrm{d}u\,\mathrm{d}v
\end{aligned} \tag{S69}$$

using

$$\begin{aligned}
f_{\tau_R|\tau_{S_0}}(v|u) &= \frac{\mathrm{d}}{\mathrm{d}u}\mathbb{P}[\tau_R \le v|\tau_{S_0} = u] \\
&= \frac{\mathrm{d}}{\mathrm{d}u}\mathbb{P}[D \le v-u|\tau_{S_0} = u] \\
&\xrightarrow{g/k\searrow 0} \frac{\mathrm{d}}{\mathrm{d}u}\mathbb{P}[D \le v-u] \\
&= f_D(v-u).
\end{aligned} \tag{S70}$$

Thus, in this limiting regime, we can split the integral with respect to $t$ by

$$\begin{aligned}
\mathbb{E}[Z_t^m] &= \int_0^\infty \int_0^\infty \mathbb{E}[Z_t^m|\tau_{S_0} = u, \tau_R = v] f_{\tau_{S_0}}(u) f_D(v-u)\,\mathrm{d}u\,\mathrm{d}v \\
&= \int_0^t \Bigg( \int_u^t \mathbb{E}[Z_t^m|\tau_{S_0} = u, \tau_R = v] f_D(v-u)\,\mathrm{d}v \\
&\qquad + \int_t^\infty \mathbb{E}[Z_t^m|\tau_{S_0} = u, \tau_R = v] f_D(v-u)\,\mathrm{d}v \Bigg) f_{\tau_{S_0}}(u)\,\mathrm{d}u \\
&\quad + \int_t^\infty \Bigg( \int_t^u \mathbb{E}[Z_t^m|\tau_{S_0} = u, \tau_R = v] f_D(v-u)\,\mathrm{d}v \\
&\qquad + \int_u^\infty \mathbb{E}[Z_t^m|\tau_{S_0} = u, \tau_R = v] f_D(v-u)\,\mathrm{d}v \Bigg) f_{\tau_{S_0}}(u)\,\mathrm{d}u.
\end{aligned} \tag{S71}$$

Label each integral by $I_A$ to $I_D$ sequentially, and we observe the following:

($I_A$) The integral goes from $\tau_{S_0} = u \in [0,t)$ and $\tau_R = v \in [u,t)$, i.e., $0 \le \tau_{S_0} \le \tau_R < t$. This means we are observing at a time $t$ after the initial depletion of both $S_0$ and $S_R$, so we are in the third phase.

($I_B$) The integral goes from $\tau_{S_0} = u \in [0,t)$ and $\tau_R = v \in [t,\infty)$, i.e., $0 < \tau_{S_0} < t \le \tau_R$. This means we are observing at a time after the initial depletion of $S_0$ but before that of $R$, so we are in the second phase.

($I_C$) The integral goes from $\tau_{S_0} = u \in [t,\infty)$ and $\tau_R = v \in [t,u)$, i.e., $t \le \tau_R < \tau_{S_0} < \infty$. As $\tau_{S_0} \le \tau_R$ with probability converging to one, the integral is supported on a measure-zero set, and evaluates to zero.

($I_D$) The integral goes from $\tau_{S_0} = u \in [t,\infty)$ and $\tau_R = v \in [u,\infty)$, i.e., $t \le \tau_{S_0} \le \tau_R < \infty$. This means we are observing at a time before the initial depletion of both $S_0$ and $R$, so we are in the first phase.

Denote $I^{(\mathrm{C,i})} = I_D, I^{(\mathrm{C,ii})} = I_B, I^{(\mathrm{C,iii})} = I_A$ and $I_C = 0$, then we have

$$I^{(\mathrm{C,i})} = \mathbb{E}[Z_t^m \mathbb{1}_{\{\tau_{S_0} \geq t\}} \mathbb{1}_{\{\tau_R \geq t\}}], \tag{S72}$$

$$I^{(\mathrm{C,ii})} = \mathbb{E}[Z_t^m \mathbb{1}_{\{\tau_{S_0} < t\}} \mathbb{1}_{\{\tau_R \geq t\}}], \tag{S73}$$

$$I^{(\mathrm{C,iii})} = \mathbb{E}[Z_t^m \mathbb{1}_{\{\tau_{S_0} < t\}} \mathbb{1}_{\{\tau_R < t\}}]. \tag{S74}$$

In the (C, i) phase, as $X_t = \alpha - \beta + Y_t$ and $Z_t = \beta - Y_t$, we have

$$\begin{aligned} f_{X_t,Y_t}(x,y) &= \mathbb{P}[X_t = x, Y_t = y] \\ &= \mathbb{P}[Y_t = x - \alpha + \beta, Y_t = y] \\ &= \delta_{x-\alpha+\beta,y} P_y^{(\mathrm{C,i})}(t), \end{aligned} \tag{S75}$$

so

$$\begin{aligned} \mathbb{E}^{(\mathrm{C,i})}[Z_t^n] &= \int_0^\infty \int_0^\infty (\beta - y)^n f_{X_t,Y_t}(x,y)\,\mathrm{d}x\,\mathrm{d}y \\ &= \int_0^\infty \int_0^\infty (\beta - y)^n \delta_{x-\alpha+\beta,y} P_y^{(\mathrm{C,i})}(t)\,\mathrm{d}\mu_{\mathbb{Z}}(x)\,\mathrm{d}\mu_{\mathbb{Z}}(y) \\ &= \sum_{y=\beta-\alpha+1}^{\beta} (\beta - y)^n P_y^{(\mathrm{C,i})}(t). \end{aligned} \tag{S76}$$

In the (C, ii) phase, as $Z_t = (\alpha + \beta - Y_t)/2$, so

$$\begin{aligned} \mathbb{E}^{(\mathrm{C,ii})}[Z_t^n] &= \int_0^t \mathbb{E}\left[\left(\frac{\alpha+\beta-Y_t}{2}\right)^n \mathbb{1}_{\{Y_t>0\}} \middle| \tau_{S_0} = u\right] f_{\tau_{S_0}}(u)\,\mathrm{d}u \\ &= \int_0^t \mathbb{E}\left[\left(\frac{\alpha+\beta-Y_{t-u}}{2}\right)^n \mathbb{1}_{\{Y_{t-u}>0\}}\right] f_{\tau_{S_0}}(u)\,\mathrm{d}u \\ &= \frac{1}{2^n}\int_0^t \left[\sum_{y\in\mathcal{Y}^{(\mathrm{C,ii})}\setminus\{0\}} (\alpha+\beta-y)^n P_y^{(\mathrm{C,ii})}(t-u)\right] f_{\tau_{S_0}}(u)\,\mathrm{d}u, \end{aligned} \tag{S77}$$

where

$$\begin{aligned} f_{\tau_{S_0}}(u) &= \frac{\mathrm{d}}{\mathrm{d}u}\mathbb{P}[X_u = 0] \\ &= \frac{\mathrm{d}}{\mathrm{d}u}\mathbb{P}[Y_u = \beta - \alpha] \\ &= \sum_{n=\beta-\alpha+1}^{\beta} \kappa_n^{(\mathrm{C,i})} e^{-\lambda_n^{(\mathrm{C,i})} u}. \end{aligned} \tag{S78}$$

In the (C, iii) phase,

$$\begin{aligned} \mathbb{E}^{(\mathrm{C,iii})}[Z_t^n] &= \int_0^t \int_u^t \mathbb{E}[Z_t^m | \tau_{S_0} = u, \tau_R = v] f_{\tau_{S_0}}(u) f_D(v-u)\,\mathrm{d}v\,\mathrm{d}u \\ &= \int_0^t \int_0^v \mathbb{E}[Z_t^n | \tau_{S_0} = u, \tau_R = v] f_{\tau_{S_0}}(u) f_D(v-u)\,\mathrm{d}u\,\mathrm{d}v \\ &= \int_0^t \mathbb{E}[Z_t^n | \tau_R = v] \left(\int_0^v f_{\tau_{S_0}}(u) f_D(v-u)\,\mathrm{d}u\right)\mathrm{d}v \\ &= \int_0^t \mathbb{E}[Z_t^n | \tau_R = v] f_{\tau_R}(v)\,\mathrm{d}v \\ &= \int_0^t \sum_{z=0}^{\lfloor(\alpha+\beta)/2\rfloor} z^n P_z^{(\mathrm{C,iii})}(t-v) f_{\tau_R}(v)\,\mathrm{d}v, \end{aligned} \tag{S79}$$

where

$$f_{\tau_R}(v) = \frac{\mathrm{d}}{\mathrm{d}v}\mathbb{P}[Y_v = 0]$$
$$= \begin{cases} \sum_{n\in\mathscr{Y}_2^{(\mathrm{C,ii})}} \kappa_n^{(\mathrm{C,ii})} e^{-\lambda_n^{(\mathrm{C,ii})} v}, & 2 \mid (\beta-\alpha), \\ \sum_{n\in\mathscr{Y}_1^{(\mathrm{C,ii})}} \kappa_n^{(\mathrm{C,ii})} e^{-\lambda_n^{(\mathrm{C,ii})} v}, & 2 \nmid (\beta-\alpha). \end{cases} \tag{S80}$$

Accordingly, we have

$$\mathbb{E}^{(\mathrm{C})}[X_t^n] = \sum_{y=\beta-\alpha+1}^{\infty} (\alpha-\beta+y)^n P_y^{(\mathrm{C,i})}(t) + \int_0^t \left[\sum_{z=0}^{\infty} (\alpha+\beta-2z)^n P_z^{(\mathrm{C,iii})}(t-v)\right] f_{\tau_R}(v)\,\mathrm{d}v, \tag{S81}$$

$$\mathbb{E}^{(\mathrm{C})}[Y_t^n] = \sum_{y=\beta-\alpha+1}^{\infty} y^n P_y^{(\mathrm{C,i})}(t) + \int_0^t \left(\sum_{y\in\mathscr{Y}^{(\mathrm{C,ii})}\setminus\{0\}} y^n P_y^{(\mathrm{C,ii})}(t-u)\right) f_{\tau_{S_0}}(u)\,\mathrm{d}u, \tag{S82}$$

$$\mathbb{E}^{(\mathrm{C})}[Z_t^n] = \sum_{y=\beta-\alpha+1}^{\infty} (\beta-y)^n P_y^{(\mathrm{C,i})}(t) + \frac{1}{2^n}\int_0^t \left[\sum_{y\in\mathscr{Y}^{(\mathrm{C,ii})}\setminus\{0\}} (\alpha+\beta-y)^n P_y^{(\mathrm{C,ii})}(t-u)\right] f_{\tau_{S_0}}(u)\,\mathrm{d}u$$
$$+ \int_0^t \left(\sum_{z=0}^{\infty} z^n P_z^{(\mathrm{C,iii})}(t-v)\right) f_{\tau_R}(v)\,\mathrm{d}v. \tag{S83}$$

Using (S54), we can then write

$$\int_0^t \sum_{y\in\mathscr{Y}^{(\mathrm{C,ii})}\setminus\{0\}} (L(y))^n P_y^{(\mathrm{C,ii})}(t-u) f_{\tau_{S_0}}(u)\,\mathrm{d}u$$
$$= \sum_{y\in\mathscr{Y}^{(\mathrm{C,ii})}\setminus\{0\}} \frac{(L(y))^n}{\lambda_y^{(\mathrm{C,ii})}} \sum_{q\in\mathscr{Y}_y^{(\mathrm{C,ii})}} \sum_{r=\beta-\alpha+1}^{\beta} \kappa_q^{(\mathrm{C,ii})}\kappa_r^{(\mathrm{C,i})} h_t(\lambda_q^{(\mathrm{C,ii})}, \lambda_r^{(\mathrm{C,i})}), \tag{S84}$$

and

$$\int_0^t \sum_{z=0}^{\lfloor(\alpha+\beta)/2\rfloor} (L(z))^n P_z^{(\mathrm{C,ii})}(t-v) f_{\tau_R}(v)\,\mathrm{d}v$$
$$= \sum_{z=0}^{\lfloor(\alpha+\beta)/2\rfloor} (L(z))^n \binom{\lfloor(\alpha+\beta)/2\rfloor}{z} \sum_{q=0}^{\lfloor(\alpha+\beta)/2\rfloor - z} \sum_{r\in\mathscr{Y}_\sigma^{(\mathrm{C,ii})}} \binom{\lfloor(\alpha+\beta)/2\rfloor - z}{q} (-1)^q \kappa_r^{(\mathrm{C,ii})} h_t\left(g(z+q), \lambda_r^{(\mathrm{C,ii})}\right), \tag{S85}$$

where

$$\sigma := 1 + \mathbb{1}_{\{2\mid(\beta-\alpha)\}}. \tag{S86}$$

### S8 Solutions to the Mean-Field Ordinary Differential Equations

Recall the system of ordinary differential equations

$$\frac{\mathrm{d}c_{S_0}}{\mathrm{d}t} = -\bar{k}c_{S_0}c_R + 2\bar{g}c_{S_R}, \tag{S87}$$

$$\frac{\mathrm{d}c_R}{\mathrm{d}t} = -\bar{k}c_{S_0}c_R, \tag{S88}$$

$$\frac{\mathrm{d}c_{S_R}}{\mathrm{d}t} = \bar{k}c_{S_0}c_R - \bar{g}c_{S_R}, \tag{S89}$$

and

$$\frac{\mathrm{d}c_S}{\mathrm{d}t} = \bar{g}c_{S_R}. \tag{S90}$$

#### S8.1 The Fast-Growing Limit ($\bar{g}/\bar{k}_0 \nearrow \infty$)

For $\bar{g}/\bar{k}_0 \nearrow \infty$, consider the quasi-steady-state approximation

$$\frac{\mathrm{d}c_{S_R}^{(\mathrm{A})}}{\mathrm{d}t} = 0. \tag{S91}$$

This leads to

$$c_{S_R}^{(\mathrm{A})}(t) = \frac{\bar{k}}{\bar{g}}c_{S_0}^{(\mathrm{A})}(t)c_R^{(\mathrm{A})}(t) \xrightarrow{g/k_0 \nearrow \infty} 0. \tag{S92}$$

Substituting (S92) into (S87) yields

$$\frac{\mathrm{d}c_{S_0}^{(\mathrm{A})}}{\mathrm{d}t} = \bar{k}c_{S_0}^{(\mathrm{A})}c_R^{(\mathrm{A})}, \tag{S93}$$

such that the system of equations can be reduced to

$$\frac{\mathrm{d}c_{S_0}^{(\mathrm{A})}}{\mathrm{d}t} = \bar{\alpha} + \bar{\beta} - \frac{\mathrm{d}c_R^{(\mathrm{A})}}{\mathrm{d}t}, \tag{S94}$$

$$\frac{\mathrm{d}c_R^{(\mathrm{A})}}{\mathrm{d}t} = -kc_R^{(\mathrm{A})}(\bar{\alpha} + \bar{\beta} - c_R^{(\mathrm{A})}), \tag{S95}$$

$$\frac{\mathrm{d}c_{S_R}^{(\mathrm{A})}}{\mathrm{d}t} = 0, \tag{S96}$$

using (S2), which have the solutions

$$c_{S_0}^{(\mathrm{A})}(t) = \bar{\alpha} + \bar{\beta} - \frac{\bar{\alpha} + \bar{\beta}}{(\bar{\alpha}/\bar{\beta})e^{(\bar{\alpha}+\bar{\beta})\bar{k}t} + 1} \tag{S97}$$

$$c_R^{(\mathrm{A})}(t) = \frac{\bar{\alpha} + \bar{\beta}}{(\bar{\alpha}/\bar{\beta})e^{(\bar{\alpha}+\bar{\beta})\bar{k}t} + 1}, \tag{S98}$$

$$c_{S_R}^{(\mathrm{A})}(t) = 0. \tag{S99}$$

#### S8.2 The Resource-Limited Fast-Feeding Limit ($\bar{g}/\bar{k}_0 \searrow 0, \bar{\alpha} \geq \bar{\beta}$)

For $\bar{g}/\bar{k}_0 \searrow 0$ and $\bar{\alpha} \geq \bar{\beta}$, take

$$\lim_{t \searrow 0} c_{S_0}^{(\mathrm{B,ii})}(t) = \bar{\alpha} - \bar{\beta}, \tag{S100}$$

$$\lim_{t \searrow 0} c_R^{(\mathrm{B,ii})}(t) = 0, \tag{S101}$$

$$\lim_{t \searrow 0} c_{S_R}^{(\mathrm{B,ii})}(t) = \bar{\beta}, \tag{S102}$$

which serve as the boundary conditions between (B,i) and (B,ii) in Figure 2. Note that it is not feasible to analytically consider the initial rapid depletion of $R$ in the deterministic case due to the inherent smoothness of the differential equation model [30].

After $R$ is all consumed, only growth occurs, such that for $t > 0$, we have the equations

$$\frac{\mathrm{d}c_{S_0}^{(\mathrm{B,ii})}}{\mathrm{d}t} = 2\bar{g}c_{S_0}^{(\mathrm{B,ii})}, \tag{S103}$$
$$\frac{\mathrm{d}c_{R}^{(\mathrm{B,ii})}}{\mathrm{d}t} = 0, \tag{S104}$$
$$\frac{\mathrm{d}c_{S_R}^{(\mathrm{B,ii})}}{\mathrm{d}t} = -\bar{g}c_{S_0}^{(\mathrm{B,ii})}, \tag{S105}$$

which have the solutions

$$c_{S_0}^{(\mathrm{B,ii})}(t) = \bar{\alpha} + \bar{\beta}(1 - 2e^{-\bar{g}t}), \tag{S106}$$
$$c_{R}^{(\mathrm{B,ii})}(t) = 0, \tag{S107}$$
$$c_{S_R}^{(\mathrm{B,ii})}(t) = \bar{\beta}e^{-\bar{g}t}. \tag{S108}$$

So, overall, we have

$$c_{S_0}^{(\mathrm{B})}(t) = \begin{cases} \bar{\alpha}, & t = 0, \\ \bar{\alpha} + \bar{\beta}(1 - 2e^{-\bar{g}t}), & t > 0, \end{cases} \tag{S109}$$
$$c_{R}^{(\mathrm{B})}(t) = \begin{cases} \bar{\beta}, & t = 0, \\ 0, & t > 0, \end{cases} \tag{S110}$$
$$c_{S_R}^{(\mathrm{B})}(t) = \begin{cases} 0, & t = 0, \\ \bar{\beta}e^{-\bar{g}t}, & t > 0. \end{cases} \tag{S111}$$

### S8.3 The Species-Limited Fast-Feeding Limit ($\bar{g}/\bar{k}_0 \searrow 0, \bar{\alpha} < \bar{\beta}$)

For $\bar{g}/\bar{k}_0 \searrow 0$ and $\bar{\alpha} < \bar{\beta}$, take

$$\lim_{t \searrow 0} c_{S_0}^{(\mathrm{C,ii})}(t) = 0, \tag{S112}$$
$$\lim_{t \searrow 0} c_{R}^{(\mathrm{C,ii})}(t) = \bar{\beta} - \bar{\alpha}, \tag{S113}$$
$$\lim_{t \searrow 0} c_{S_R}^{(\mathrm{C,ii})}(t) = \bar{\alpha}. \tag{S114}$$

Also, consider the quasi-steady-state approximation

$$\frac{\mathrm{d}c_{S_0}^{(\mathrm{C,ii})}}{\mathrm{d}t} = 0, \tag{S115}$$

which only holds for $t \in (0, \bar{\tau})$ such that $c_{S_0}^{(\mathrm{C,ii})}(t) \geq 0$ on the interval.

This leads to

$$c_{S_0}^{(\mathrm{C,ii})}(t) = \frac{2\bar{g}}{\bar{k}} \left( \frac{c_{S_R}^{(\mathrm{C,ii})}(t)}{c_{R}^{(\mathrm{C,ii})}(t)} \right) \xrightarrow{\bar{g}/\bar{k}_0 \searrow 0} 0. \tag{S116}$$

Substituting (S116) into (S88) yields

$$\frac{\mathrm{d}c_{R}^{(\mathrm{C,ii})}}{\mathrm{d}t} = 2\bar{g}c_{S_R}^{(\mathrm{C,ii})}, \tag{S117}$$

such that the system of equations can be reduced to

$$\frac{\mathrm{d}c_{S_0}^{(\mathrm{C,ii})}}{\mathrm{d}t} = 0, \tag{S118}$$

$$\frac{\mathrm{d}c_{R}^{(\mathrm{C,ii})}}{\mathrm{d}t} = -g(\bar{\alpha} + \bar{\beta} - c_{R}^{(\mathrm{C,ii})}), \tag{S119}$$

$$\frac{\mathrm{d}c_{S_R}^{(\mathrm{C,ii})}}{\mathrm{d}t} = -\frac{1}{2}\frac{\mathrm{d}c_{R}^{(\mathrm{C,ii})}}{\mathrm{d}t}, \tag{S120}$$

using (S2), which have the solutions

$$c_{S_0}^{(\mathrm{C,ii})}(t) = 0, \tag{S121}$$

$$c_{R}^{(\mathrm{C,ii})}(t) = \bar{\alpha}(1 - 2e^{\bar{g}t}) + \bar{\beta}, \tag{S122}$$

$$c_{S_R}^{(\mathrm{C,ii})}(t) = \bar{\alpha}e^{\bar{g}t}. \tag{S123}$$

This allows us to find $\bar{\tau}$ by setting $c_{R}^{(\mathrm{C,ii})}(\bar{\tau}) = 0$:

$$\bar{\tau} = \frac{1}{\bar{g}} \ln\left(\frac{1}{2}\left(1 + \frac{\bar{\beta}}{\bar{\alpha}}\right)\right). \tag{S124}$$

To ensure piecewise continuity at this boundary, we need to consider

$$c_{S_R}^{(\mathrm{C,ii})}(\bar{\tau}) = c_{S_R}^{(\mathrm{C,iii})}(\bar{\tau}) = \frac{1}{2}(\bar{\alpha} + \bar{\beta}). \tag{S125}$$

Also, because $c_R(t) = 0$ for $t \geq \bar{\tau}$, we have the system of equations

$$\frac{\mathrm{d}c_{S_0}^{(\mathrm{C,iii})}}{\mathrm{d}t} = -2\frac{\mathrm{d}c_{S_R}^{(\mathrm{C,iii})}}{\mathrm{d}t}, \tag{S126}$$

$$\frac{\mathrm{d}c_{R}^{(\mathrm{C,iii})}}{\mathrm{d}t} = 0, \tag{S127}$$

$$\frac{\mathrm{d}c_{S_R}^{(\mathrm{C,iii})}}{\mathrm{d}t} = -\bar{g}c_{S_R}, \tag{S128}$$

using (S2), which have the solutions

$$c_{S_0}^{(\mathrm{C,iii})}(t) = (\bar{\alpha} + \bar{\beta})\left[1 - \frac{1}{2\bar{\alpha}}(\bar{\alpha} + \bar{\beta})e^{-\bar{g}t}\right], \tag{S129}$$

$$c_{R}^{(\mathrm{C,iii})}(t) = 0, \tag{S130}$$

$$c_{S_R}^{(\mathrm{C,iii})}(t) = \frac{1}{4\bar{\alpha}}(\bar{\alpha} + \bar{\beta})^2 e^{-\bar{g}t}, \tag{S131}$$

for $t \geq \bar{\tau}$.

So, overall, we have

$$c_{S_0}^{(\mathrm{C})}(t) = \begin{cases} \bar{\alpha}, & t = 0, \\ 0, & t \in (0, \bar{\tau}), \\ (\bar{\alpha} + \bar{\beta}) \left[ 1 - \dfrac{1}{2\bar{\alpha}} (\bar{\alpha} + \bar{\beta}) e^{-\bar{g}t} \right], & t \in [\bar{\tau}, \infty), \end{cases} \tag{S132}$$

$$c_{R}^{(\mathrm{C})}(t) = \begin{cases} \bar{\beta}, & t = 0, \\ \bar{\alpha}(1 - 2e^{\bar{g}t}) + \bar{\beta}, & t \in (0, \bar{\tau}), \\ 0, & t \in [\bar{\tau}, \infty), \end{cases} \tag{S133}$$

$$c_{S_R}^{(\mathrm{C})}(t) = \begin{cases} 0, & t = 0, \\ \bar{\alpha} e^{\bar{g}t}, & t \in (0, \bar{\tau}), \\ \dfrac{1}{4\bar{\alpha}} (\bar{\alpha} + \bar{\beta})^2 e^{-\bar{g}t}, & t \in [\bar{\tau}, \infty). \end{cases} \tag{S134}$$

It is important to note that these analytical approximations to the actual numerical solutions of the system of ordinary differential equations are piecewise functions which are not differentiable at $t = 0$ and $t = \bar{\tau}$ by construction. As certain quantities are forced to be zero at some regimes, these analytical approximations do not suffer from the atto-fox problem [30].

### S9 Supplementary Figures for Results and Discussion

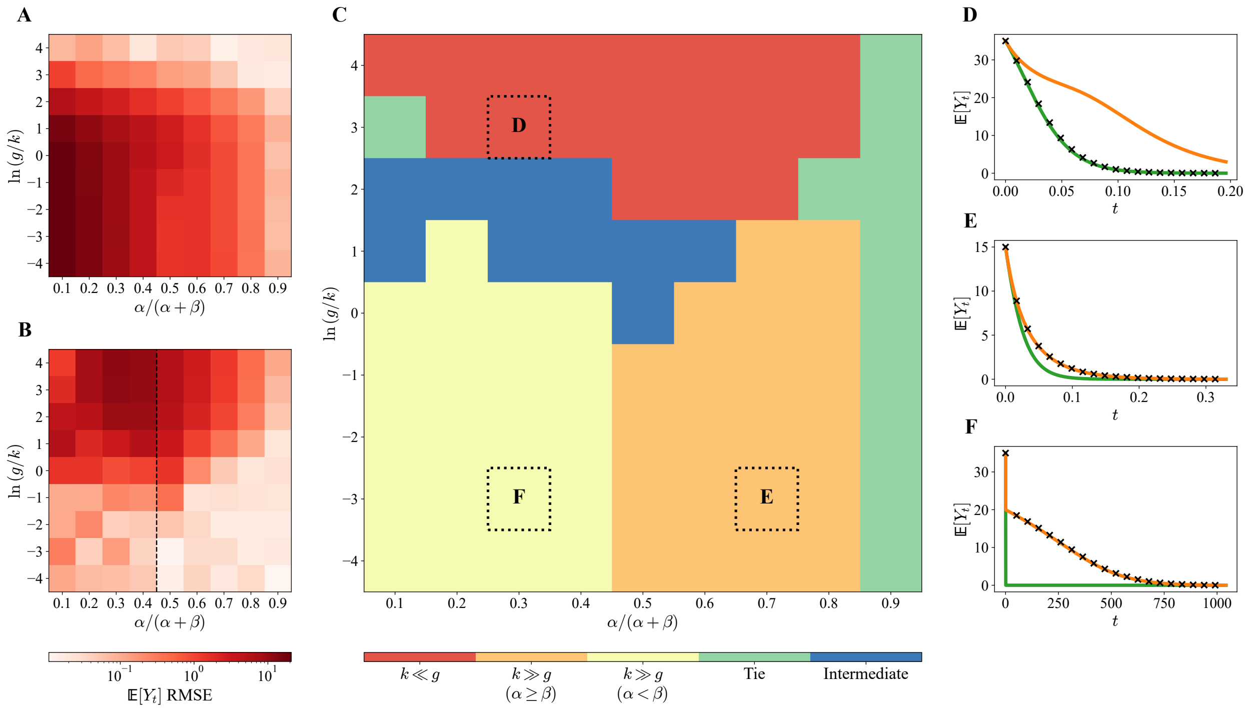

Figure S9.1: Limiting analytical solutions to the coarse-grained forward master equations are valid descriptions of the system's behaviors under relaxed values of $k$ and $g$. (**A**) Root-mean-square error (RMSE) between the fast-growing analytical approximation of $\mathbb{E}[Y_t]$ and its simulated values. (**B**) RMSE between the fast-feeding analytical approximation of $\mathbb{E}[Y_t]$ and its simulated values. The dotted black line partitions the heatmap into a $\alpha \geq \beta$ portion and a $\alpha < \beta$ portion. (**C**) Classification of single-species-single-resource resource consumption dynamics in the $\alpha/(\alpha+\beta)$-$\ln(g/k)$ plane into distinct functional forms. (**D**) Resource consumption dynamics in the relaxed fast-growing regime $(\alpha/(\alpha+\beta) = 0.3, \ln(g/k) = 3)$. (**E**) Resource consumption dynamics in the relaxed resource-limited fast-feeding regime $(\alpha/(\alpha+\beta) = 0.7, \ln(g/k) = -3)$. (**F**) Resource consumption dynamics in the relaxed species-limited fast-feeding regime $(\alpha/(\alpha+\beta) = 0.7, \ln(g/k) = -3)$. Orange and green curves correspond to analytical approximations of $\mathbb{E}[Y_t]$ in the limiting regimes $g/k \searrow 0$ and $g/k \nearrow 0$ respectively. Black crosses correspond to the simulated mean. $k$ and $\alpha+\beta$ are fixed at 1 and 50 respectively.

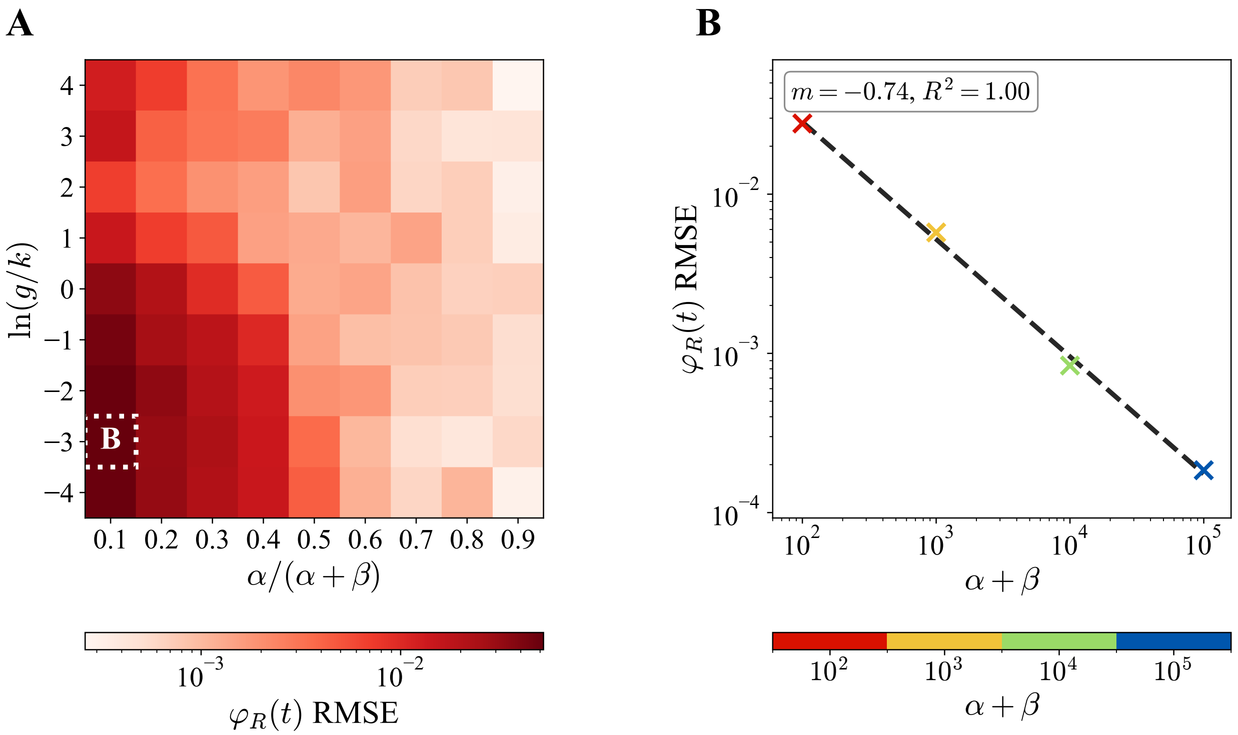

Figure S9.2: Deviation between the simulated solutions from the stochastic and deterministic frameworks. (**A**) Root-mean-square error (RMSE) between the stochastically simulated and deterministic $\varphi_R(t)$ curves. (**B**) Scaling between the $\varphi_R(t)$ RMSE with system size at the parameter point with maximum RMSE. The dotted line is the best-fit line. $k$ is fixed at 1.

### S10 Derivation of the Monod-Like Growth Curves

Recall the definition of specific growth rate [1, 31], which is

$$\mu := \frac{\mathrm{d}\ln c_S}{\mathrm{d}t} = \frac{1}{c_S}\frac{\mathrm{d}c_S}{\mathrm{d}t}. \tag{S135}$$

Substituting (S90) into (S135), we have

$$\mu = \frac{\bar{g}c_{S_R}}{c_{S_0} + c_{S_R}}. \tag{S136}$$

In the fast-feeding limit, we can plug (S92) into (S136) to get

$$\begin{aligned}\mu_{k \ll g} &= \frac{\bar{g}(\bar{k}/\bar{g})c_{S_0}c_R}{c_{S_0} + (\bar{k}/\bar{g})c_{S_0}c_R} \\ &= \frac{\bar{g}c_R}{(\bar{g}/\bar{k}) + c_R}.\end{aligned} \tag{S137}$$

In the fast-feeding limit, consider the typical case where $\bar{\alpha} < \bar{\beta}$, then we can plug (S116) into (S136) to get

$$\begin{aligned}\mu_{k \gg g} &= \frac{\bar{g}c_{S_R}}{(2\bar{g}/\bar{k})(c_{S_R}/c_R) + c_{S_R}} \\ &= \frac{\bar{g}c_R}{(2\bar{g}/\bar{k}) + c_R}.\end{aligned} \tag{S138}$$

Note that in reality, the terms in (S92) and (S116) tend to zero but are not exactly zero, such that the factor cancellations above are mathematically valid.
